# Alternative splicing of *clock* transcript mediates the response of circadian clocks to temperature changes

**DOI:** 10.1101/2024.05.10.593646

**Authors:** Yao D. Cai, Gary K. Chow, Sergio Hidalgo, Xianhui Liu, Kiya C. Jackson, Cameron D. Vasquez, Zita Y. Gao, Vu H. Lam, Christine A. Tabuloc, Haiyan Zheng, Caifeng Zhao, Joanna C. Chiu

## Abstract

Circadian clocks respond to temperature changes over the calendar year, allowing organisms to adjust their daily biological rhythms to optimize health and fitness. In *Drosophila*, seasonal adaptations and temperature compensation are regulated by temperature-sensitive alternative splicing (AS) of *period* (*per*) and *timeless* (*tim*) genes that encode key transcriptional repressors of clock gene expression. Although *clock* (*clk*) gene encodes the critical activator of clock gene expression, AS of its transcripts and its potential role in temperature regulation of clock function have not been explored. We therefore sought to investigate whether *clk* exhibits AS in response to temperature and the functional changes of the differentially spliced transcripts. We observed that *clk* transcripts indeed undergo temperature-sensitive AS. Specifically, cold temperature leads to the production of an alternative *clk* transcript, hereinafter termed *clk*-cold, which encodes a CLK isoform with an in-frame deletion of four amino acids proximal to the DNA binding domain. Notably, serine 13 (S13), which we found to be a CK1α-dependent phosphorylation site, is among the four amino acids deleted in CLK-cold protein. Using a combination of transgenic fly, tissue culture, and *in vitro* experiments, we demonstrated that upon phosphorylation at CLK(S13), CLK-DNA interaction is reduced, thus decreasing CLK occupancy at clock gene promoters. This is in agreement with our findings that CLK occupancy at clock genes and transcriptional output are elevated at cold temperature, which can be explained by the higher amounts of CLK-cold isoforms that lack S13 residue. This study provides new insights into the complex collaboration between AS and phospho-regulation in shaping temperature responses of the circadian clock.

## Introduction

Circadian clocks regulate daily physiological and behavioral rhythms to optimize health and fitness in organisms from all domains of life^1–7^. In animals and other eukaryotes, the pace of the circadian clock is highly responsive to environmental cues, with light-dark cycles as the strongest and most-studied time cue to entrain and reset circadian clocks^8^. Temperature is another important environmental cue that greatly affects the physiology of animals. Although circadian clocks can maintain its pace over a wide range of temperatures, a phenomenon called temperature compensation^9^, circadian clocks can also adapt to more extreme temperatures. For example, under cold conditions, the expression of *Period2*, a key component in the molecular clock, is induced in brown adipose tissue of mice to increase systemic heat production and resistance to cold temperature^10^. At present, there are still significant knowledge gaps regarding the molecular mechanisms by which circadian clocks respond and adapt to cold temperatures to optimize daily biological rhythms.

A number of studies have established alternative splicing (AS) as an important mechanism that mediate temperature responses of circadian clocks in animals^11–13^, plants^14^, and fungi^15–17^. The molecular clocks in most organisms are composed of transcription-translation feedback loops (TTFLs)^2,5^ that regulate daily rhythmicity in clock gene expression. The TTFL model in animal clocks was first formulated in *Drosophila* in 1990^18^. It is composed of positive elements, CLOCK (CLK) and CYCLE (CYC), that drive the transcription of clock genes including genes encoding the negative elements, PERIOD (PER) and TIMELESS (TIM), which feedback to repress the activity of positive elements. The next daily cycle of transcription begins as repression is relieved by proteasomal degradation of PER and TIM.

There has been great progress in understanding the temperature-sensitive AS of negative elements in *Drosophila* clocks, in parallel to that in fungal^15–17^ and plant clocks^14,19,20^. AS of *per* and *tim* improves fitness of flies by minimizing locomotor activity during the time with adverse environmental temperature, such as cold night and hot midday^13,21,22^. Cold-induced AS of *per dmpi8* intron at the 3’ UTR region^13^ promotes advanced evening locomotor activity peak by stimulating transcription of *daywake*^23^. In addition to *per*, four splice variants of *tim* are found to be differentially expressed in response to temperature changes^21,24,25^. Constitutively spliced *tim*-L (full-length isoform) is dominant at moderate temperature (25°C)^25^. In colder temperatures (10-18°C), increased expression of *tim*-cold and *tim*-short and cold (*tim*-sc) are linked to advanced evening locomotor activity peak. Increased expression of *tim*-M in warmer temperatures (29-35°C) contributes to the delay of evening behavioral peak^21^. However, whether transcripts of the positive elements in *Drosophila* clocks undergo temperature-sensitive AS remains unexplored. This is surprising given CLK, in partner with CYC (homolog of mammalian BMAL1), are the major transcriptional activators of clock gene expression and are critical for driving clock output^26,27^.

In comparison to the paucity of studies that explore the regulation of CLK function by AS, there have been experimental evidence and mathematical modeling that demonstrated the importance of posttranslational regulation of CLK activity. In particular, phosphorylation plays key roles in shaping the daily oscillation of CLK transcriptional activity, while CLK protein abundance stays constant over the 24-hour day-night cycle^28–30^. In midday to early evening, newly synthesized hypophosphorylated CLK binds DNA and exhibits high transcriptional activity after nuclear entry^28,31^. Due to PER-dependent recruitment of kinases, highly phosphorylated CLK appearing at late night exhibits downswing of transcriptional activity and is gradually removed from circadian promoters^28,32,33^. Dephosphorylation of CLK that occurs in early day replenishes the pool of transcriptionally active hypophosphorylated CLK^30^. However, it is unknown how the daily CLK phosphorylation profile and transcriptional activity change with temperature. Furthermore, whether post-transcriptional mechanism such as AS modulates daily CLK phosphorylation profile to regulate cold adaptation of molecular clocks remains a significant gap in knowledge.

Here we observed that cold temperature increases the usage of an alternative 3’ splice site in *clk* exon 2, leading to the production of an alternative transcript that encodes a CLK protein missing four amino acids proximal to the N-terminal DNA basic helix-loop-helix (bHLH) binding domain. We named this shorter transcript hereinafter *clk*-cold. We demonstrated that CLK-cold protein exhibits increased occupancy at CLK target gene promoters and elevates CLK target gene expression at cold temperature. Among the four amino acids deleted in CLK-cold protein, we used mass spectrometry proteomics and phosphospecific antibodies to show that CLK(S13) is phosphorylated by casein kinase 1α (CK1α). By analyzing the behavioral and molecular output of *clk*(S13D) phosphomimetic mutants, we provided evidence supporting the function of CLK(S13) phosphorylation in modulating CLK occupancy at CLK target gene promoters. This maintains daily CLK target mRNA expression and PDF neuropeptide accumulation rhythm to drive rhythmic locomotor activities. Based upon these findings, we propose a model describing a mechanism by which AS of *clk* transcript mediates the response of circadian clocks to temperature changes by modulating transcriptional activity of CLK via CK1α-dependent phosphorylation.

## Results

### Temperature regulates alternative 3’ splice site selection in exon 2 of *clk*

We first sought to determine whether *clk* transcripts exhibit alternative splicing, as in the case of other key clock genes such as *per* and *tim*. When *Drosophila clk* transcripts were first cloned in 1998 by three independent labs, two different cDNA products were identified^34–36^. The longer transcript, which we termed *clk-*long, encodes the canonical CLK protein. The other slightly shorter transcript, hereinafter termed *clk*-cold, encodes a CLK protein with a four amino acid (aa) deletion at aa13-16 (Fig. 1A). The annotation of the genomic *clk* sequence revealed a potential alternative 3’ splice site in exon 2 of *clk*, which may have produced the two cDNA of different lengths. Indeed, both transcripts are expressed in fly heads, according to deep sequencing of circadian transcriptome in Hughes et al^37^. Cell-specific RNA-seq data from Wang et al.^38^ also indicated that both *clk-*long and *clk-*cold are expressed in several circadian neuronal cell types, including small lateral ventral neurons (sLN_v_), the key pacemaker neurons (Fig. S1A). Given that the four aa deletion of CLK-cold is adjacent to the bHLH DNA-binding domain (aa17-62)^35,36^, we hypothesize that differential expression of these two transcripts could impact the function of the molecular clock.

**Fig. 1.**
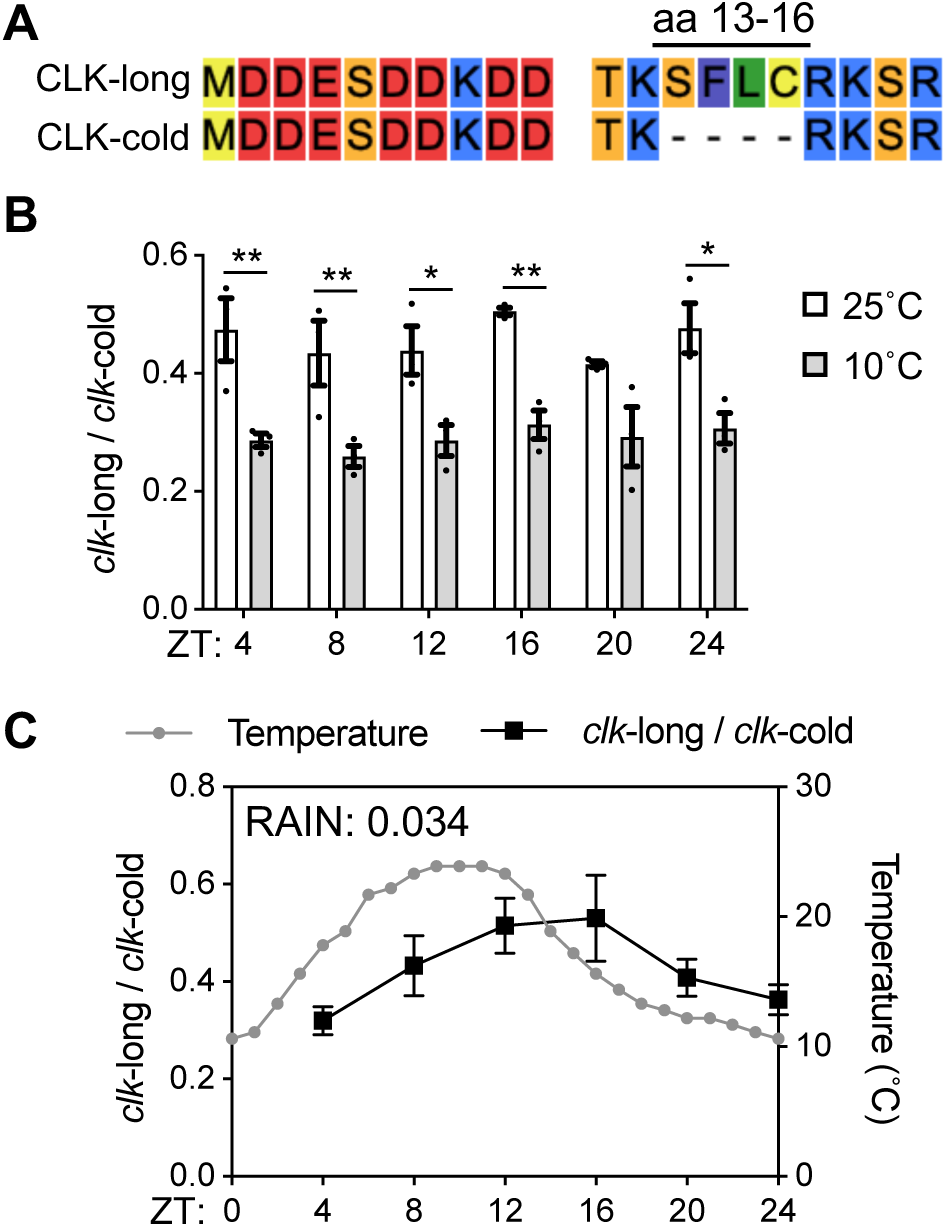
Cold induces alternative splicing of *clk*. **A.** Alignment of amino acid sequences encoded by two *clk* transcripts isolated from heads of *w*^1118^ flies. Four amino acids (aa 13-16) are absent in CLK-cold. **B-C.** The ratio of *clk*-long to *clk*-cold was measured in heads of *w*^1118^ flies by quantitative RT-PCR. Flies were entrained in 12h:12h LD at indicated constant temperature in **B** and 14h:10h LD at natural daily temperature cycles in **C**. Flies were collected on LD3 at indicated time-points (ZT). Error bars indicate ± SEM (n=3), **p<0.01, *p<0.05, Two-Way ANOVA and Šídák’s post hoc test.

To further confirm the expression of both *clk-*long and *clk-*cold, we generated *clk* cDNA fragments that span exons 1 and 2 from RNA extracted from whole heads of *w*^1118^ flies collected at a time-point when *clk* mRNA expression is high^35^ (Fig. S1B). We noticed three potentially different cDNAs when we analyzed the amplified cDNAs on agarose gel (Fig. S1C). Sanger sequencing revealed that the top band represents *clk*-long, while the bottom two bands represent *clk*-cold (Fig. S1D).

To determine whether AS of *clk* is temperature-sensitive, we evaluated daily relative abundance of *clk*-long and *clk*-cold from head extracts of *w*^1118^ flies entrained under LD at 25°C and 10°C, respectively (Fig. 1B). 10°C was chosen to better simulate a more naturalistic cold temperature, e.g. morning of a spring day, at which *Drosophila* flies are viable and active^39^. Nested qPCR assays targeting the alternative 3’ splice site region allows quantitative analysis of both transcripts (Fig. S1E). At 25°C, *clk-* long and *clk-*cold were expressed at 1:2 ratio. However, the relative abundance of *clk*-long decreases and *clk*-cold becomes the dominant isoform at 10°C. AS of *clk* did not exhibit daily oscillation at constant temperature, given the ratio does not oscillate under LD condition, as determined by rhythmicity analysis incorporating nonparametric methods (RAIN)^40^ (25°C: p=0.81; 10°C: p=0.86, RAIN).

Even in endothermic organisms such as mice, body temperature rhythm was found to drive rhythmic AS of over 1000 exons^12^. We therefore hypothesize that environmental temperature cycles drive rhythmic AS of *clk* in ectothermic *Drosophila*. We measured expression of *clk-*long and *clk-*cold under semi-natural conditions, where the incubators were set to mimic a typical day in May in Davis, California, USA (weatherspark.com), with diurnal temperatures ranging from 10°C to 25°C (Fig. 1C). We observed steady increase of relative *clk*-long in response to increasing temperature in the first half of the day. As temperature decreases in the second half of the day, relative level of *clk*-long decreases. Under environmental temperature cycles, AS of *clk* is rhythmic (p=0.034, RAIN). These data suggest that AS of *clk* is sensitive to environmental temperature changes.

### Elevated CLK-DNA binding contributes to increased CLK target mRNA level at cold temperature

Since CLK-cold is missing four amino acids (aa 13-16) adjacent to the N-terminal bHLH DNA binding domain (aa17-62), we hypothesize that CLK-cold displays altered CLK-DNA binding activity. To test this hypothesis, we performed CLK chromatin immunoprecipitation (CLK-ChIP) followed by qPCR using extracts from adult fly heads collected at 25°C vs 10°C. In agreement with published data^41^, we observed rhythmic CLK occupancy at *per* CRS, a region within the promoter critical for generating rhythmic *per* expression^42^ and *vri E-box* (CLK binding motif) at 25°C (Fig. 2A and B) (*per* CRS: p=0.014; *vri E-box*: p=0.002, RAIN). However, we observed dampening of daily rhythmicity of CLK occupancy at 10°C (*per* CRS: p=0.582; *vri E-box*: p=0.142, RAIN), most likely due to significantly higher CLK occupancy at both circadian promoters at ZT4. Our ChIP data suggest CLK-cold exhibits elevated DNA binding activity as compared to CLK-long, most notably at ZT4.

**Fig. 2.**
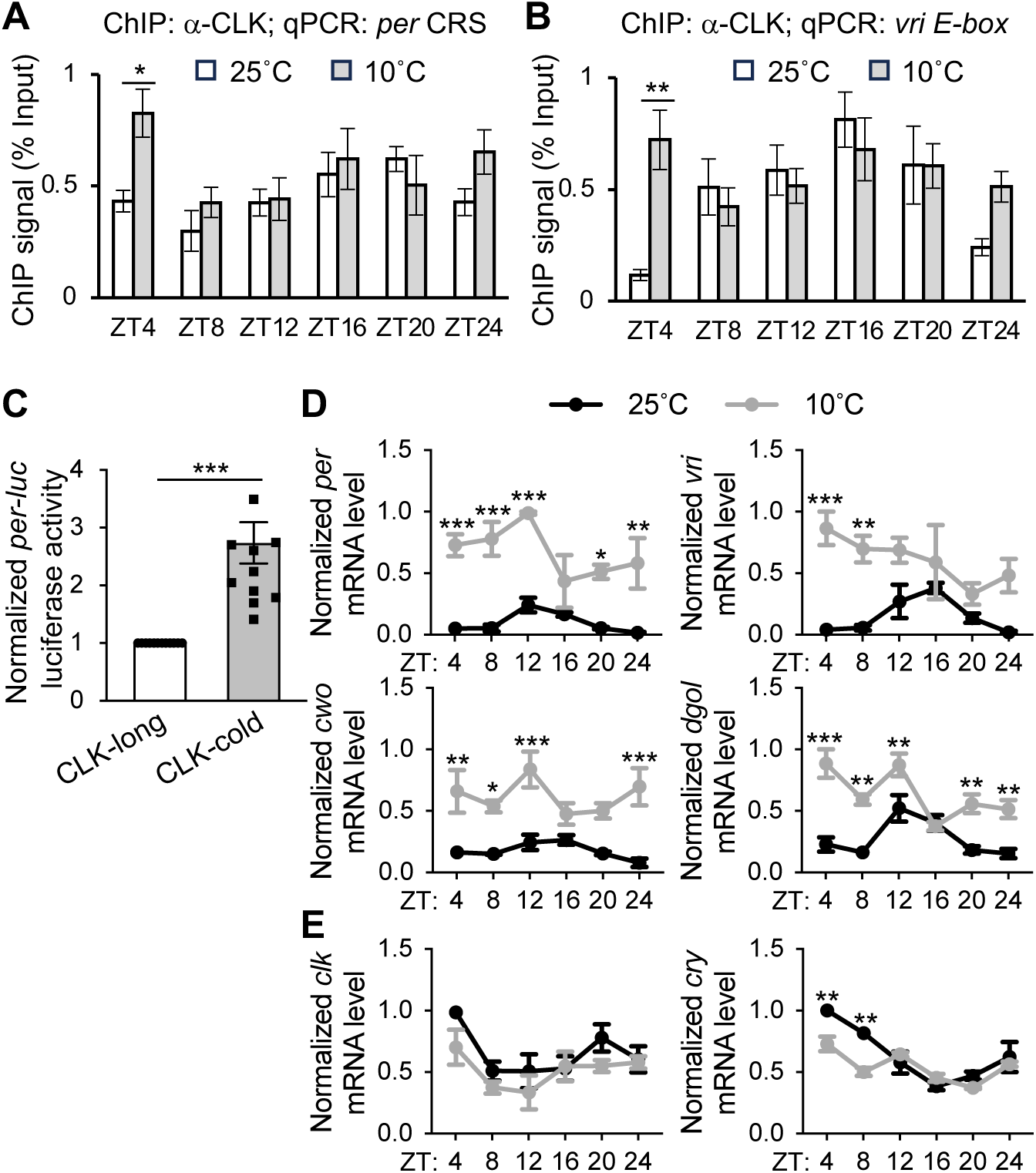
Elevated CLK-DNA binding contributes to increased mRNA level of CLK targets at low temperature. **A-B.** ChIP assays using fly head extracts comparing daily CLK occupancy at *per* and *vri* promoter in *w*^1118^ flies collected at 25°C and 10°C. CLK-ChIP signals were normalized to % input. ChIP signals for an intergenic region were used for non-specific background deduction. Flies were entrained in 12h:12h LD and collected on LD3 at indicated time-points (ZT) (n=5). Error bars indicate ± SEM, **p< 0.01, *p< 0.05, Two-Way ANOVA and Šídák’s post hoc test. **C.** *per-E-box-luciferase* (*per-luc*) reporter assay performed in *Drosophila* S2 cells. Luciferase activity was normalized to CLK-long and expressed as fold change relative to CLK-long. Error bars indicate ± SEM (n=12), ***p<0.001, two-tailed Student’s t test. **D-E.** Steady state daily mRNA expression of CLK targets (*per*, *vri*, *cwo* and *dgol*) and non-CLK targets (*clk* and *cry*) in heads of *w*^1118^ flies. Flies were entrained in 12h:12h LD and collected on LD3 at indicated temperatures and time-points (ZT) (n=3). Error bars indicate ± SEM, ***p<0.001, **p< 0.01, *p< 0.05, Two-Way ANOVA and Šídák’s post hoc test.

We next asked whether altered DNA binding activity regulates transcriptional activity of each CLK isoforms. We first assayed transcriptional activity of CLK in *Drosophila* S2 cells using a *per*-*luc* reporter assay^36^. We compared *per-luc* reporter gene activity in S2 cells expressing CLK-long or CLK-cold (Fig. 2C). The 2.7-fold increased reporter gene activity in CLK-cold suggests a significantly elevated transcriptional activity of CLK-cold, as compared to CLK-long. Transcriptional activity of CLK-cold was also inferred by measuring expression of known CLK target genes in flies entrained in 12h:12h LD at 25°C vs 10°C, respectively (Fig. 2D). At 10°C where CLK-cold is dominant, mRNA levels of CLK targets, including *per*, *vrille (vri)*, *clockwork orange (cwo)*, *goliath* (*dgol)*, are significantly higher than the levels observed at 25°C at multiple time-points. Similarly, total *tim* and *pdp1* mRNA levels are elevated at 10°C, despite the fact that these two genes have temperature-sensitive AS (Fig. S2). In contrast, mRNA levels of non-CLK targets including *clk* and *cryptochrome* (*cry*) did not increase at 10°C (Fig. 2E). This strongly indicates that the observed elevated expression at cold temperature is not a general phenomenon. Taken together, our results revealed that cold-induced AS of *clk* results in a CLK protein that binds more readily to DNA at certain times of the day-night cycle, thereby increasing expression of CLK target genes.

### CLK(S13) is a CK1α-dependent phosphorylation site that regulates transcriptional activity of CLK

We next sought to explain how the four aa deletion in CLK-cold alters CLK transcriptional output. Since CLK phosphorylation status displays daily rhythmicity and is tightly correlated with its transcriptional activity^28,30,43^, we investigated whether these four amino acids overlap with any kinase-targeted motif. Kinase prediction using group-based phosphorylation site predicting and scoring (GPS) 5.0^44^ showed that the serine 13 (S13) residue could be a potential substrate of kinases from several kinase families (Table S1). Previous studies suggested that NEMO^45^, SGG^46–48^, CK2^49^ and CK1α^50^ could be CLK kinases and CLK kinases could be recruited by PER to phosphorylate CLK. Among these kinases, GPS 5.0 kinase prediction algorithm identified CK1α to be the most likely candidate to phosphorylate CLK(S13).

To first determine if phosphorylation of S13 can regulate CLK transcriptional activity, we generated *clk* plasmids expressing non-phosphorylatable S13 to Alanine (A) or phosphomimetic S13 to Aspartic Acid (D) mutations, both in the context of in CLK-long isoform, and tested their transcriptional activity using *per-luc* luciferase assays in *Drosophila* S2 cells. Whereas CLK(S13A) was found to exhibit higher transcriptional activity when compared to CLK-long (WT), CLK(S13D) had significantly lower transcriptional activity (Fig. 3A). Our results suggest S13 could indeed be a potential phosphorylation site that regulates CLK transcriptional activity. We then performed a series of experiments to determine if CLK(S13) is a *bona fide* CK1α-dependent phosphorylation site. We first determined if CLK interacts with CK1α by performing coimmunoprecipitation (coIP) assays using protein extracts from *Drosophila* S2 cells coexpressing CLK-V5 and CK1α-cmyc (Fig. 3B-D). We detected interactions between CLK and CK1α when using CLK-V5 as bait (Fig. 3B and C). Reciprocal coIP using CK1α-cmyc as bait resulted in the same conclusion (Fig. 3B and D). Control experiments were performed using extracts of S2 cells expressing either of the proteins alone to demonstrate minimal non-specific binding (Fig. 3B-D).

**Fig. 3.**
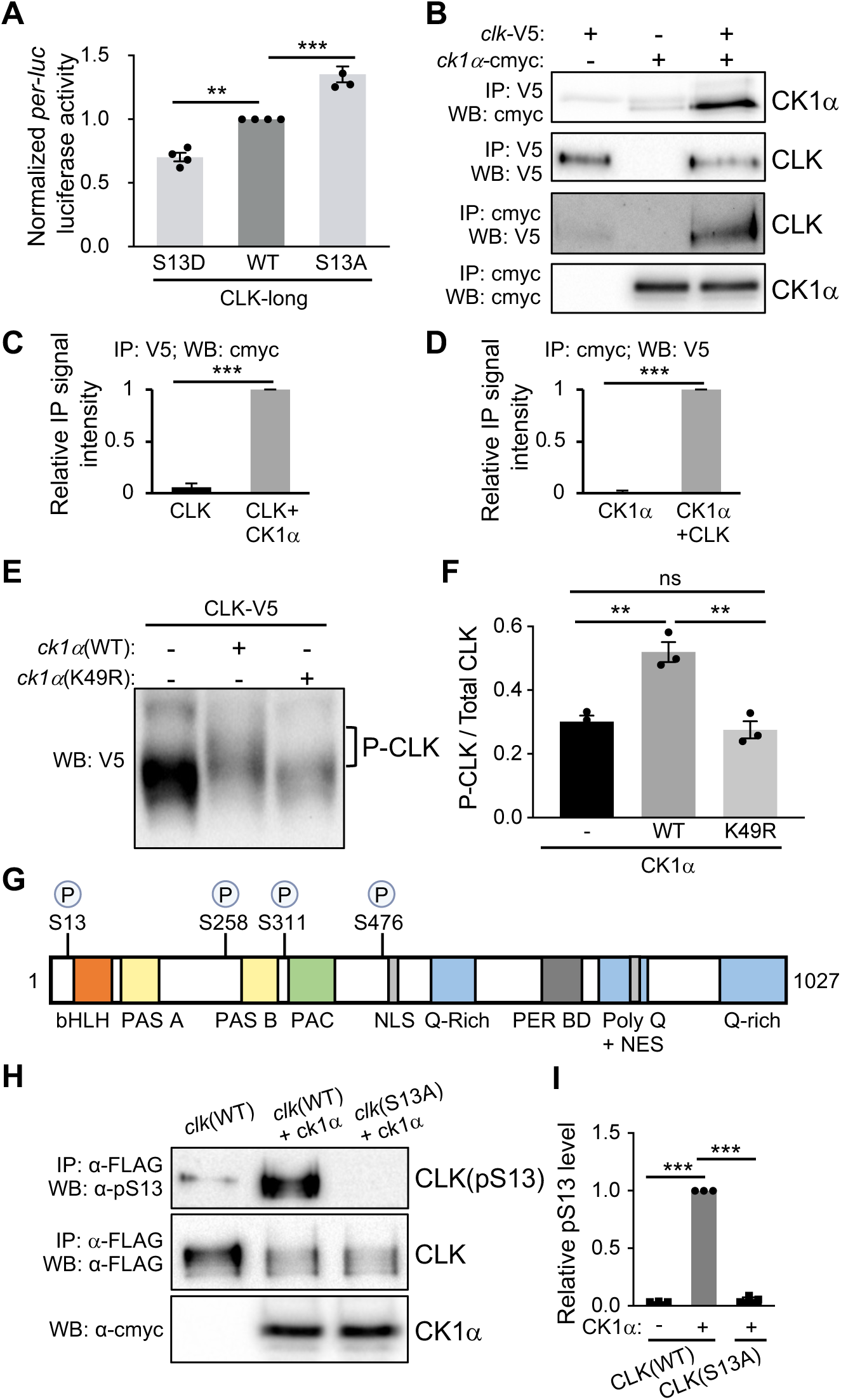
CLK(S13) is a substrate of CK1α. **A.** *per-E-box-luciferase* (*per-luc*) reporter assay performed in *Drosophila* S2 cells. Luciferase activity was normalized to CLK(WT) and expressed as fold change relative to CLK-long. Error bars indicate ± SEM (n=4), ***p<0.001, One-Way ANOVA and Dunnett post hoc test. **B.** Western blots showing reciprocal coimmunoprecipitations (coIPs) to examine the interactions of CLK and CK1α. S2 cells were cotransfected with 0.8μg of pAc-*clk*-V5-His and 0.8μg of pMT-*ck1α*-6xc-myc or transfected with a single plasmid for control experiments. Protein extracts were divided into two equal aliquots, and each aliquot was independently incubated with either α-c-myc beads or α-V5 beads. Immuno-complexes were analyzed by western blotting in the presence of the indicated antibody. **C-D.** Bar graphs displaying quantification of reciprocal coIPs (**B**). Values for binding are normalized to amount of bait detected in the IPs and expressed as relative signal intensity (maximum value=1). Error bars indicate ± SEM (n=3), two-tailed Student’s t test. **E** Western blots showing mobility shift of CLK on a Phos-tag SDS-PAGE. S2 cells were transfected with 0.8μg of pAc-*clk*-V5 together with 0.6μg of either pMT-*ck1α*-FH, pMT-*ck1*α(K49R)-FH, or pMT-FH. **F.** Quantification of phosphorylated/total CLK in **E**. Error bars indicate ± SEM (n=3). ***p<0.001, **p<0.01, One-Way ANOVA and Tukey’s post hoc test. **G.** Schematic diagram depicting *Drosophila melanogaster* CLK (amino acid 1-1027) adapted from Mahesh et al., 2014 and CK1α-dependent phosphorylation sites identified by mass spectrometry. Previously described domains of CLK: basic helix-loop-helix (bHLH) (aa 17-62)^35,36^; PAS-A (aa 96-144)^35,36^; PAS B (aa 264-309)^35,36^; C-terminal of PAS domain (PAC) (aa 315-379)^35^; NLS (aa 480-494)^31^; PER binding domain (PER BD) (aa 657-707)^107^; Q-rich regions (aa 546-575, aa 957-1027), Poly-Q (aa 552-976)^35,36^ and NES (aa 840-864)^31^. **H.** *Drosophila* S2 cells were transfected with pAc-*clk*(WT)-FLAG or pAc-*clk*(S13A)-FLAG and cotransfected with an empty plasmid (pMT-cmyc-His) or pMT-*ck1*α-cmyc. Protein extracts were incubated with α-FLAG resin. Total CLK isoforms, CLK(pS13), and CK1α protein levels were analyzed by Western Blotting with the indicated antibodies. **I.** Bar graph showing relative CLK pS13 levels in **H** normalized to total CLK isoforms. Error bars indicate ± SEM (n=3), ***p<0.001, One-Way ANOVA and Dunnett post hoc test.

Next, we determined if CLK is phosphorylated by CK1α by assessing CLK mobility shift on SDS-PAGE. We analyzed CLK in protein extracts of *Drosophila* S2 cells expressing either CLK alone or CLK coexpressed with CK1α. We observed slower-migrating CLK isoforms on a regular SDS-PAGE gel (Fig. S3), likely representing phosphorylated CLK. Phos-Tag SDS-PAGE gel^51^ was used to enhance phosphorylation-dependent mobility shift (Fig. 3E and F). In addition, to test whether CK1α catalytic activity is responsible for the observed mobility shift, we coexpressed CLK with either CK1α(WT) or CK1α(K49R), a kinase-dead variant^50^. We observed substantial slower-migrating CLK species in the presence of CK1α(WT). The amount of slower-migrating species was significantly reduced with CK1α(K49R) coexpression. These results indicate that CK1α kinase activity is required for CLK mobility shift.

To specifically determine whether CLK(S13) is phosphorylated by CK1α, we leveraged mass spectrometry (MS) to identify CK1α-dependent phosphorylation sites of CLK expressed in *Drosophila* S2 cells. This cell culture system has previously been used to map physiologically relevant phosphorylation sites on *Drosophila* PER^47,52,53^, TIM^54^ and CLK^55,56^. We coexpressed CLK tagged with FLAG epitope with either CK1α(WT) or kinase dead CK1α(K49R) in S2 cells and performed FLAG affinity purifications prior to MS analysis. We identified eight phosphorylation sites on CLK (Table S2 and Lam^57^). Among them, we identified four sites that exhibited elevated phosphopeptide abundance when coexpressed with CK1α(WT) as compared to CK1α(K49R) (Fig. 3G and Lam^57^). These CK1α-dependent sites include S13, which is next to the bHLH DNA binding domain^35,36^; S258 and S311 next to PAS B protein binding domain^35,36^; S476 next to the nuclear localization signal (NLS)^31^.

Finally, to further validate the phosphorylation of S13, we generated a S13 phosphospecific antibody (α-pS13) and assayed CLK(S13) phosphorylation in protein extracts of *Drosophila* S2 cells expressing *clk*-V5 (WT) with or without *ck1α* (Fig. 3H and I). Immunoblotting showed that CLK(pS13) significantly increased when *ck1α* was coexpressed (Fig. 3H, lanes 1 and 2). Importantly, there was little to no α-pS13 signal detected in extracts of S2 cells coexpressing *clk*(S13A) and *ck1α* (Fig. 3H, lane 3, top panel), suggesting that α-pS13 antibody is phosphospecific. Taken together, our results strongly support that CLK(S13) is a CK1α*-*dependent phosphorylation site.

### Flies harboring mutations at CLK(S13) display altered circadian behavioral rhythms

So far, our results suggest that S13 phosphorylation reduces CLK transcriptional activity. Furthermore, we show that temperature-sensitive AS at cold temperature led to increased production of CLK-cold that lacks this inhibitory S13 phosphorylation, thus promoting CLK target mRNA expression in the cold. To characterize the function of CLK(S13) phosphorylation *in vivo*, we generated transgenic fly lines expressing non-phosphorylatable CLK(S13A) or phosphomimetic CLK(S13D) variants. These mutated *clk* genes are expressed under endogenous *clk* promotor^55^. p{*clk*(X)-V5} (X represents WT, S13A or S13D variants) transgenic fly lines were crossed into *clk*^out^ background^55^ to remove endogenous *clk* expression. Next, we monitored daily locomotor activity rhythms of *clk* transgenic flies, given it is a robust behavioral output of the circadian clock^58^ (Fig. 4A). Flies were entrained for 4 days in 12h:12h Light:Dark (LD) cycles followed by release in 7 days in constant darkness (DD) to assess free-running rhythms. As expected, *clk*^out^ null mutant exhibited arrhythmic locomotor activity in DD, similar to published results^55^. *clk*^out^ flies expressing a *clk*(WT) transgene displayed robust daily activity rhythms with a ∼24-hour period, indicating effective rescue of the arrhythmic *clk*^out^ mutation (Fig. 4A and Table S3). As compared to *clk*(WT), *clk*(S13D) exhibited reduced rhythmicity and period lengthening by 1.8 hours. *clk*(S13A) also displayed period lengthening by 1.5 hours and reduced rhythmicity as compared to *clk*(WT) flies. Taken together, our data suggest that CLK(S13) phosphorylation is required for robust circadian timekeeping.

**Fig. 4.**
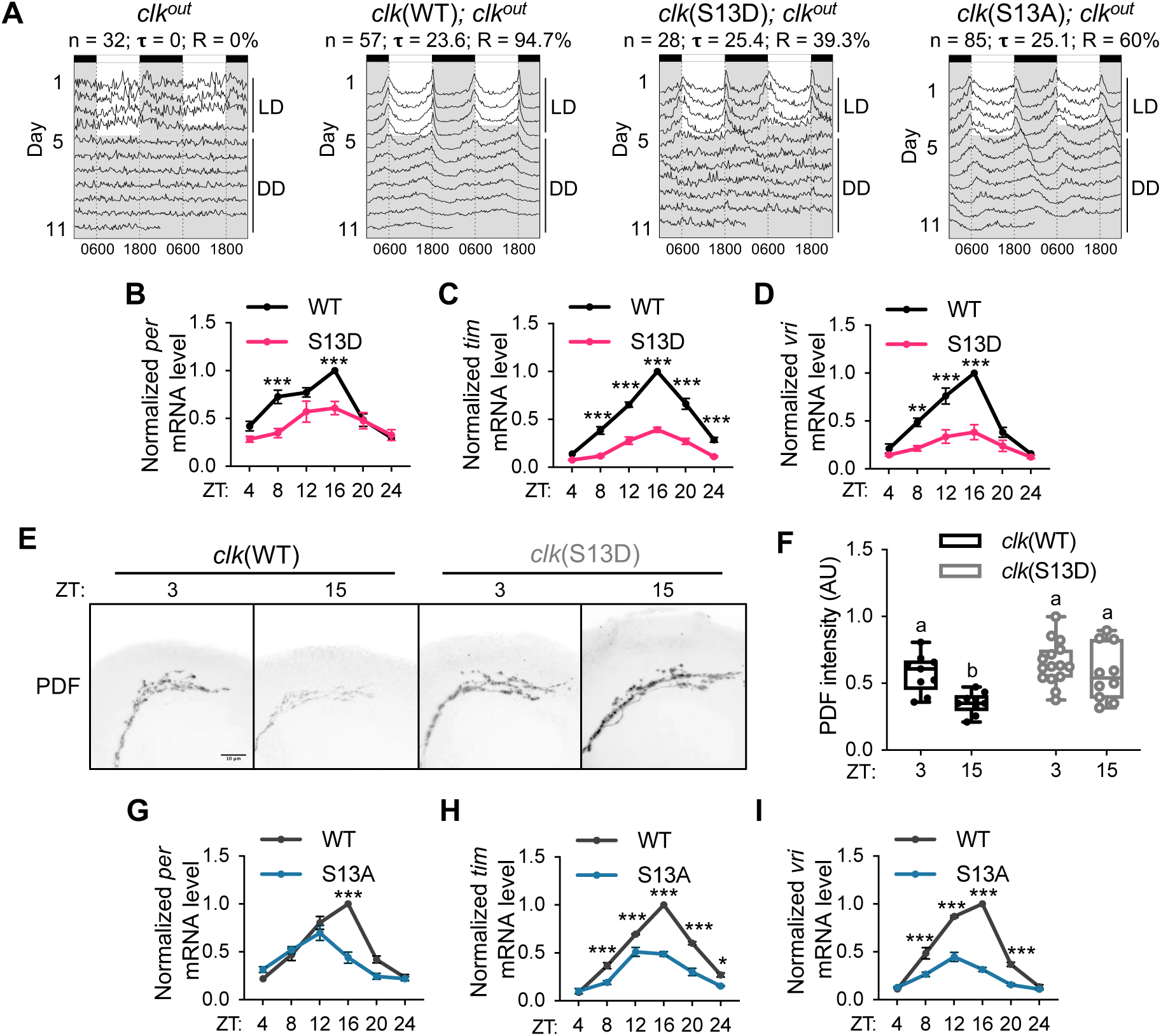
Flies expressing CLK(S13) variants display altered circadian behavioral and molecular output. **A.** Double-plotted actograms of flies harboring transgenes for altered S13, a CK1α-dependent phosphorylation site. Average activity of each genotype was plotted using FaasX. n represents the sample size; Tau (ι−) represents the average period length of the indicated group of flies in constant darkness (DD). R represents percentage of flies that are rhythmic. Flies were entrained for 4 days in 12h:12h LD and then left in DD for 7 days. **B-D.** Steady state daily mRNA expression of CLK targets (*per*, *tim*, and *vri*) in heads of *clk*(WT); *clk*^out^ and *clk*(S13D); *clk*^out^ flies. Flies were entrained in 12h:12h LD cycles at 25°C and collected on LD3 at indicated time-points (ZT) (n=3). Error bars indicate ± SEM, ***p<0.001, **p< 0.01, *p< 0.05, Two-Way ANOVA and Šídák’s post hoc test. **E.** Representative confocal images of dorsal projection of sLN_v_s neurons in adult fly brains stained with α-PDF (C7). Scale bar (merged image in *clk*(WT) ZT3) represents 10μm. Flies were entrained for 4 days in 12h:12h LD cycles and collected at the indicated times on LD4 for fixation and immunofluorescence analysis. **F.** Box plot showing the quantification of PDF intensity in **E**. Error bars indicate ± SEM, **p< 0.01, Two-Way ANOVA and Šídák’s post hoc test. **G-I** Steady state mRNA expression of CLK targets (*per*, *tim*, *vri*) in heads of *clk*(WT); *clk*^out^ and *clk*(S13A); *clk*^out^ flies. Flies were entrained in 12h:12h LD cycles at 25°C and collected on LD3 at indicated time-points (ZT) (n=3). Error bars indicate ± SEM, ***p<0.001, **p< 0.01, *p< 0.05, Two-Way ANOVA and Šídák’s post hoc test.

### CLK(S13) phosphorylation reduces CLK transcriptional activity

To determine if CLK(S13) phosphorylation-mediated downregulation in transcriptional activity observed in *Drosophila* S2 cells translates to whole animals, we quantified the mRNA of known CLK targets including *per*, *tim* and *vri* in *clk*(S13D) mutants at 25°C. We observed significant reduction in levels and cycling amplitude of all mRNAs measured in *clk*(S13D) mutants as compared to *clk*(WT) flies, as determined by CircaCompare^59^ (Fig. 4B-D and Table S4). The dampened mRNA oscillation in *clk*(S13D) flies is consistent with dampened behavioral rhythmicity (Fig. 4A). It is important to highlight that the reduced expression of CLK targets in *clk*(S13D) mutants is opposite of the higher CLK target gene expression when flies were maintained at 10°C rather than at 25°C (compare Fig. 2D and 4B-D). This suggest the phosphorylation of S13 is a key regulatory event that fails to occur when flies express CLK-cold at 10°C.

Since previous studies showed that phosphorylation-dependent reduction in CLK stability^49^ is also a plausible mechanism to reduce CLK transcriptional activity, we wanted to rule out the possibility that CK1α can modify CLK stability (Fig. S4A and B). We measured CLK degradation independent of TTFLs by performing cycloheximide (CHX) chase assays in *Drosophila* S2 cells expressing CLK in the presence or absence of CK1α. We observed similar rates of CLK protein degradation in the two conditions, suggesting that CK1α does not regulate CLK stability. In agreement with S2 cell results, we did not observe significant difference in CLK abundance between *clk*(WT) and *clk*(S13D) flies maintained in LD cycles (Fig. S4C and D).

We next sought to explore potential clock output that mediates impaired behavioral rhythmicity in *clk*(S13D) mutant. Pigment-dispersing factor (PDF), a key neuropeptide in clock neuronal circuits, has been shown as an important molecular clock output to regulate rhythmic locomotor activity in flies^60^. PDF level is modulated by a number of CLK target genes, including *vri*^61^, *hormone receptor-like 38* (HR38)^62^ and *stripe*^62^. Since we observed significant alteration of CLK target genes in *clk*(S13D) flies (Fig. 4B-D), we hypothesized altered diurnal changes of PDF level at the dorsal projection of sLN_v_ neurons may cause dampened behavioral rhythms in *clk*(S13D) mutant. We monitored PDF level using whole-mount immunocytochemistry, and observed that PDF exhibits diurnal changes in abundance in *clk*(WT) flies, which is consistent with previous studies^61,63^ (Fig. 4E and F). As expected, diurnal changes in PDF abundance are abolished in *clk*(S13D) mutant. Importantly, the constant high PDF level in *clk*(S13D) mutant is opposite of the lowered PDF level in *w*^1118^ flies at 10°C, rather than at 25°C^64^. This further supports the crucial role of S13 phosphorylation in maintaining diurnal changes of PDF level and therefore robust locomotor activity rhythms that do not occur when flies express CLK-cold at 10°C.

In addition to *clk*(S13D) flies, we also assayed CLK target gene expression in *clk*(S13A) mutants. In contrary to what we expect based on our observation of elevated transcriptional activity of CLK(S13A) variant determined by *per-luc* reporter assay in *Drosophila* S2 cells (Fig. 3A), we observed that the expression of CLK target mRNAs in *clk*(S13A) flies was also reduced when compared to *clk*(WT) (Fig. 4G-I and Table S4). This discrepancy between the extent to which the *clk*(S13A) mutation impacts CLK target gene expression in tissue culture versus in whole animals is further explored in the next section.

### CLK(S13) phosphorylation reduces CLK occupancy at CLK target gene promoters

We further investigated the molecular basis of CLK(S13) phosphorylation in regulating circadian rhythms by characterizing the impact of S13 phosphorylation in modulating CLK-DNA interactions *in vitro* and in flies. We overlayed an AlphaFold^65^ model of *Drosophila* CLK-bHLH (aa1-71 of CLK-long including the bHLH domain) to the crystal structure of human CLOCK-BMAL1-DNA^66^ and found plausible contacts between S13 and the negatively charged DNA backbone (Fig. 5A). This hints at phosphorylation being a direct modulator of CLK-DNA binding by suppressing CLK-DNA interactions via charge-charge repulsion. To test this hypothesis, we expressed and purified aa1-71 fragment of CLK-bHLH with and without the S13D mutation from *E. coli* (Fig. S5A and B) and measured their binding affinity to a 21-bp *per* promoter as bait^67^ using biolayer interferometry (BLI)^68^ (Fig. 5B). The EC_50_ between WT CLK-bHLH and DNA binding was estimated to be 0.27 μM (95% CI = [0.19, 0.37], Fig. 5C). Importantly, introduction of the phosphomimetic S13D mutation abolishes the binding of CLK-bHLH to DNA (Fig. 5C), demonstrating the potent inhibition of S13 phosphorylation on the ability of CLK to bind to DNA and act as a transcription factor. To demonstrate the specificity of the CLK-*per* promoter interaction, we performed a control experiment where the DNA is composed of a scrambled DNA sequence. We observed a reduction in CLK binding affinity on scrambled DNA compared to the 21-bp *per* promoter sequence, suggesting that the CLK-DNA binding in our BLI assay is selective for CLK target gene promotors (Fig. S5C).

**Fig. 5.**
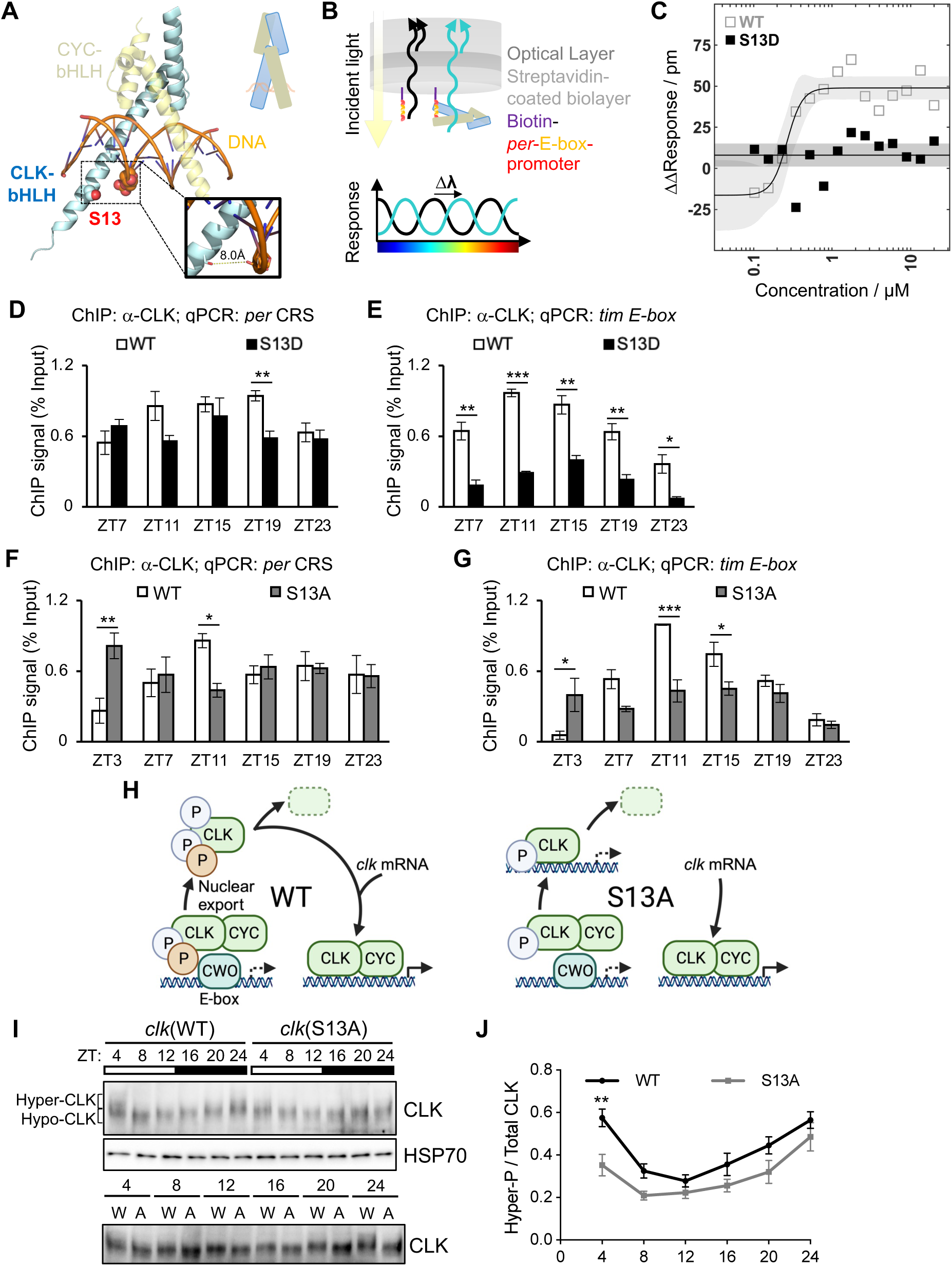
CLK(S13) phosphorylation modulates CLK-DNA binding. **A.** Model of *Drosophila* CLK (cyan)-CYC (yellow) bHLH heterodimer in complex with DNA (orange), with the side chain of S13 and the phosphate backbone of a proximal *E-box* nucleotide shown as spheres. The S13 hydroxyl oxygen atom, along with the oxygen atoms in the phosphate backbone, are shown in red. The inset shows a close-up view and the estimated distance between the two oxygen atoms. The model was generated by superimposing an AlphaFold-predicted structure of *Drosophila* CLK-bHLH (aa 1-71) with the crystal structure of human CLK and BMAL1 in complex with *E-box* DNA (PDB 4H10). A cartoon representation is shown in the upper right. **B.** Design of the Biolayer Interferometry (BLI) experiment. Biotin-labeled *per* promoter DNA was immobilized onto streptavidin-coated biosensors. CLK-bHLH-DNA interactions led to an increase in the effective thickness in the biolayer and a change in interference wavelength. **C.** Quasi-steady state signal response of CLK-bHLH-DNA binding in the presence (black, filled) and absence (grey, hollow) of the phosphomimetic S13D mutation. Solid lines and shaded areas show fits and 95% prediction interval to the 4-parameter Hill equation for CLK-bHLH-WT and nonbinding baseline for CLK-bHLH-S13D. **D-G.** ChIP assays using fly head extracts comparing CLK occupancy at *per* and *tim* promoter of indicated fly genotypes collected at 25°C on LD3 at indicated time-points (ZT) after entrainment in 12h:12h LD. CLK-ChIP signals were normalized to % input. ChIP signals for two intergenic regions were used for non-specific background deduction (**D-E**, n=3; **F-G**, n=4). Error bars indicate ± SEM, ***p<0.005, **p< 0.01, *p< 0.05, Two-Way ANOVA and Šídák’s post hoc test. **H.** Model describing the alteration of the molecular clock in flies expressing CLK(S13A) variant. In WT flies, upon CLK removal from DNA promoted by PER-dependent phosphorylation and CWO-dependent mechanisms, S13 phosphorylation prevents CLK from binding back to DNA. CLK then undergoes nuclear export and further phosphorylation^31^. Hyperphosphorylated CLK is either targeted for degradation or dephosphorylated to replenish the pool of hypophosphorylated CLK^30,31^. Dephosphorylated CLK and newly translated, hypophosphorylated CLK then promotes transcription of clock-controlled genes. In S13A flies, after initial CLK removal from DNA, increased CLK(S13A)-DNA binding affinity leads to premature binding of transcriptionally inactive CLK and DNA. This leads to reduction in CLK(S13A) nuclear export and reduce the replenishment of transcriptionally active CLK in the next cycle. PER, TIM, and phosphatase CKA-PP2A are not depicted for simplicity. Created with BioRender.com licensed to the lab of J.C. Chiu. **I.** Western blots comparing CLK protein profiles in heads of *clk*(WT); *clk*^out^ and *clk*(S13A); *clk*^out^. Flies were entrained for 4 days in 12h:12h LD and collected at the indicated times on LD3. Brackets indicate hypo- and hyperphosphorylated CLK isoforms. ⍺-HSP70 was used to indicate equal loading and for normalization. Bottom blot also detects CLK in the same two genotypes (W for WT and A for S13A) but the samples for each timepoint (ZT) were ran side by side to facilitate comparison of mobility shift. **J.** Quantification of hyperphosphorylated/total CLK. The top half of the CLK signal shown at ZT4 in *clk*(WT) flies (lane 1) is used as a reference to classify CLK isoforms as hyperphosphorylated (n=4). Error bars indicate ± SEM, **p< 0.01, Two-Way ANOVA and Šídák’s post hoc test.

We next performed CLK-ChIP followed by qPCR using extracts from adult fly heads to further determine CLK-DNA binding in fly tissues (Fig. 5D-E). We observed significantly lower CLK occupancy in *clk*(S13D) mutants as compared to *clk*(WT) flies at ZT19 at *per CRS* (Fig. 5D). CLK occupancy at *tim E-box* is also significantly lower in *clk*(S13D) mutants at all time-points tested (Fig. 5E). Together with data from *in vitro* experiments (Fig. 5A-C), our results revealed that CLK occupancy at CLK target gene promoters decreases upon CLK(S13) phosphorylation, which explains lower expression of CLK target genes in *clk*(S13D) flies.

In the case of *clk*(S13A) mutant, CLK-ChIP qPCR showed significantly higher CLK occupancy at *per CRS* and *tim E-box* at ZT3 (Fig. 5F and G). Increased CLK-DNA binding in early day in *clk*(S13A) mutant (ZT3, Fig. 5F and G) resembles our observation of higher CLK-DNA binding in WT flies at 10°C (ZT4, Fig. 2A and B). These results support our model that AS allows CLK-cold to escape inhibitory phosphorylation at S13 to promote CLK-DNA binding. Surprisingly, CLK occupancy at ZT11 (*per CRS* and *tim E-box*) and ZT15 (*tim E-box*) in *clk*(S13A) mutant is significantly lower as compared to that in *clk*(WT). How might increased CLK-DNA binding in early morning contribute to a reduction of CLK-DNA binding in the beginning of the night? Upon CLK removal from DNA, a fraction of CLK undergoes nuclear export and further phosphorylation followed by degradation^31^. Whereas another fraction of CLK is dephosphorylated to replenish the pool of hypophosphorylated CLK for the next round of CLK-activated transcription^30^. We hypothesize in *clk*(S13A) mutants, increased CLK-DNA binding affinity leads to a premature binding of transcriptionally inactive CLK and DNA in early morning. These inactive CLK proteins cannot be exported to nucleus for dephosphorylation and are eventually degraded in the nucleus. Without replenishing transcriptionally active CLK, CLK occupancy at circadian promoters in the beginning of the night is reduced (Fig. 5H). We therefore analyzed daily CLK phosphorylation in *clk*(S13A) (Fig. 5I and J). As we expected, CLK phosphorylation level in *clk*(S13A) was significantly reduced in early day (ZT3) as compared to *clk*(WT). This is likely due to increased DNA-binding activity and the inability of CLK to be exported to the cytoplasm for further phosphorylation. Notably, reduced CLK occupancy at circadian promoters when CLK target transcription peaks (ZT11-15) (Fig. 5F and G) is consistent with lower CLK target mRNA levels in *clk*(S13A) mutants (Fig. 4G-I). We cannot however rule out the possibility that additional feedback mechanisms absent in tissue culture system could have contributed to the discrepancy between reduced CLK target gene expression in *clk*(S13A) flies (Fig. 4G-I) and increased transcriptional activity of CLK(S13A) in S2 cells (Fig. 3A).

### PER-DBT interaction influences CK1α-dependent downregulation of CLK transcriptional activity

Finally, we sought to determine the molecular requirements for CK1α to phosphorylate CLK(S13). It has been proposed that PER-TIM repressor complexes recruit yet uncharacterized kinases for timely CLK phosphorylation to enhance repression^32,69^. Since CK1α has been shown to interact with PER in both the cytoplasm and the nucleus^50^, we hypothesize that PER promotes CK1α-dependent phosphorylation of CLK (Fig. 6A). We first performed *per-luc* reporter assay to measure CLK transcriptional activity in S2 cells expressing CLK and PER in the absence or presence of CK1α (Fig. 6B). As expected, PER expression downregulates CLK transcriptional activity. CK1α coexpression further reduced CLK transcriptional activity, indicating an enhanced PER repression via PER-dependent CK1α phosphorylation of CLK. We observed no significant difference between baseline luciferase activity and cells expressing CLK and PER in conjunction with CK1α, suggesting that PER and CK1α together essentially abolished transcriptional activity of CLK.

**Fig. 6.**
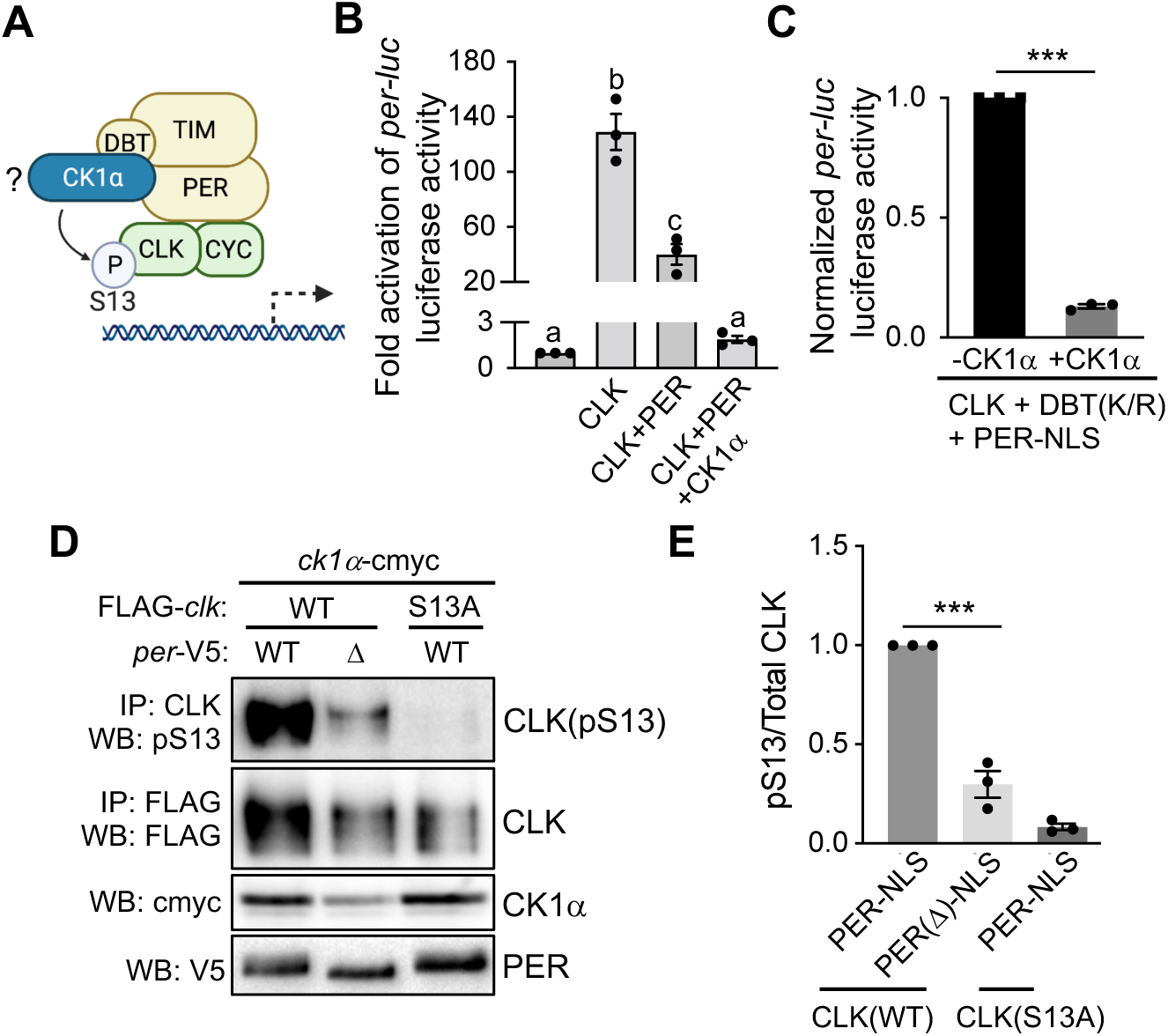
PER-DBT scaffolding promotes CK1α-dependent CLK(S13) phosphorylation. **A.** Schematic illustrating the PER-DBT scaffolding model first proposed by Yu et al.^28^. Created with BioRender.com licensed to the lab of J.C. Chiu. **B-C.** *per-E-box-luciferase* (*per-luc*) reporter assay performed in S2 cells. **B.** The fold activation of *per-luc* were graphed. Error bars indicate ± SEM (n=3). One-Way ANOVA and Tukey’s post hoc test. **C.** Luciferase activities were normalized to CLK+DBT(K/R)+PER-NLS. Error bars indicate ± SEM (n=3). **D.** *Drosophila* S2 cells were transfected with indicated plasmids. Protein extracts were incubated with α-FLAG resin. Total CLK isoforms, pCLK(S13), PER, and CK1α protein levels were analyzed by Western Blotting with indicated antibodies. **E.** Bar graph showing relative CLK pS13 levels in **D** normalized to total CLK level. Error bars indicate ± SEM (n=3), ***p<0.001, One-Way ANOVA and Šídák’s post hoc test.

CK1α regulates PER repression activity by favoring PER nuclear entry and promoting PER-DBT interaction and phosphorylation-dependent degradation^50^. To remove the confounding effect of CK1α on PER nuclear localization and degradation in our luciferase assay, we therefore expressed nuclear localization sequence (NLS)-tagged PER (PER-NLS) and DBT(K/R), a kinase-dead DBT variant^70^ to specifically examine the role of CK1α in modulating CLK activity (Fig. 6C). We observed significant reduction in CLK-dependent *per-luc* activity upon CK1α coexpression with PER-NLS and DBT(K/R). This suggests that in addition to the regulation of CK1α on PER nuclear entry and stability, the enhanced PER repression of CLK seen in Fig. 6B is also mediated through CLK phosphorylation by CK1α.

To directly determine whether PER-DBT scaffold promotes CLK(S13) phosphorylation by CK1α, we assayed CLK(S13) phosphorylation in protein extracts of *Drosophila* S2 coexpressing FLAG-*clk* and *ck1α* in combination with either *per*(WT) or *per*(Δ), a *per* mutant encoding a variant lacking PER-DBT binding domain^71^ (Fig. 6D and E). Western blotting with S13 phosphospecific antibody showed that CLK(pS13) was significantly reduced when *per*(Δ) is coexpressed, as compared to *per*(WT) (Fig. 6D, lane 1-2). As expected, little to no CLK(pS13) signal was detected in *clk*(S13A) (Fig. 6D, lane 3), showing the specificity of α-pS13. Together, our results support that PER-DBT serves as a scaffold to enhance CK1α-dependent CLK(S13) phosphorylation.

## Discussion

In this study, we report temperature-sensitive AS of the core clock gene, *clk*. Under cold condition, CLK-cold is the dominant isoform and displays higher transcriptional activity as compared to the canonical CLK-long isoform. S13 is a phosphorylation site among the four amino acids deleted in CLK-cold protein. Combining the results from a series of molecular and behavioral analyses of transgenic flies expressing non-phosphorylatable *clk*(S13A) and phosphomimetic *clk*(S13D) mutants, we formulated a model describing AS of *clk* transcripts in regulating the response of the molecular clock to temperature changes (Fig. 7). At higher temperature (25°C), CLK-long harboring S13 residue is expressed. CK1α relies on PER-DBT as a scaffold to phosphorylate CLK(S13), which prevents CLK-DNA binding in the early morning. At low temperature (10°C), dominant isoform CLK-cold lacks S13 residue, thereby escaping CK1α-dependent S13 phosphorylation. As a result, mRNA expression of CLK target genes is enhanced at low temperature.

**Fig. 7.**
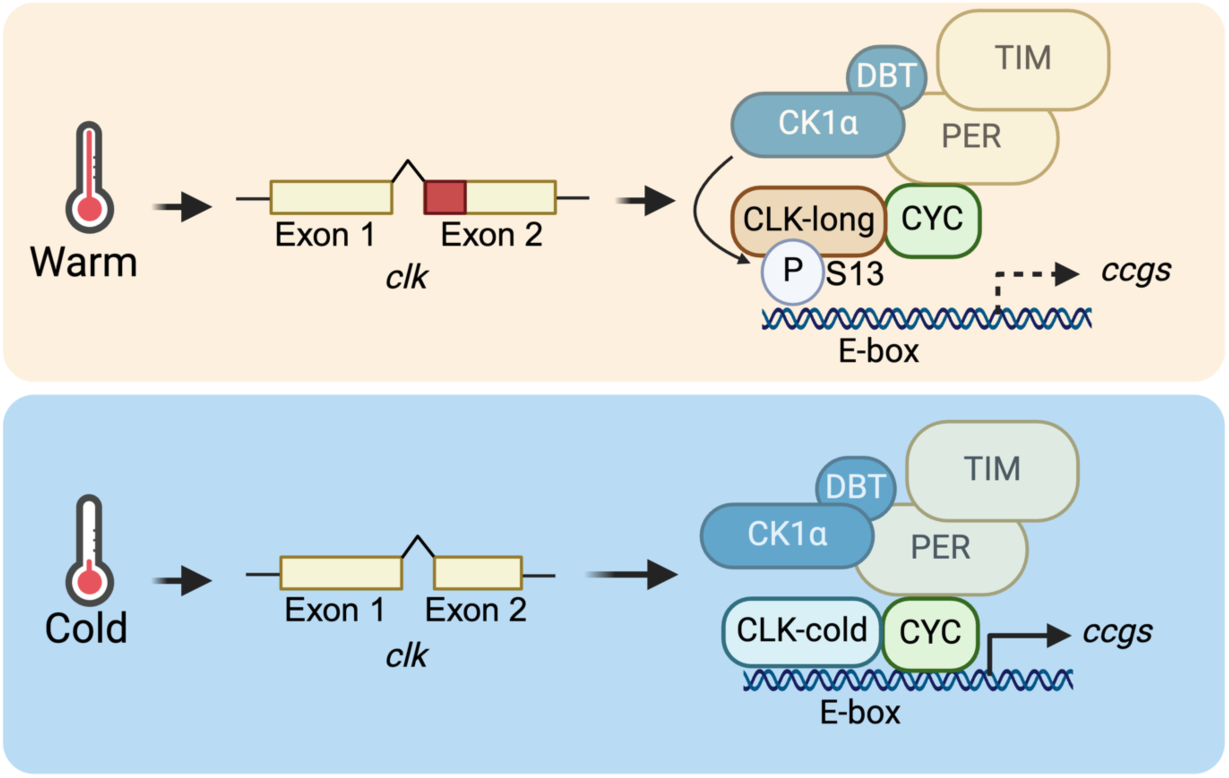
Model illustrating the regulation of the molecular clock by temperature-sensitive alternative splicing of *clk*. Top panel: At warm temperature (25°C), CLK-long isoform harboring S13 residue is expressed, due to alternative 3’ splice site selection of exon 2 of *clk*. PER-DBT scaffolding promotes CK1α-dependent phosphorylation of CLK(S13) and reduces CLK-DNA binding. Bottom panel: At cold temperature (10°C), CLK-cold isoform lacking S13 residue becomes dominant, therefore escaping this inhibitory phosphorylation. As a result, this leads to elevated mRNA expression of CLK targets under low temperature. Created with BioRender.com licensed to the lab of J.C. Chiu.

We observed daily rhythms of *clk* transcripts under environmental temperature cycles in ectothermic flies (Fig. 1). Mice and humans, despite being endothermic, display body temperature rhythm of a few degrees that is shown to entrain peripheral clocks^72,73^. Temperature-sensitive AS has been identified in over 1000 exons^12^, including genes involved in general transcription^74^ and regulating stability of core clock protein mPER1^75^. Future studies are required to fully understand the role of temperature-sensitive AS on the entrainment of peripheral clocks by body temperature rhythms. It is also interesting to point out that cold temperature also induces mRNA expression of several CLOCK target genes in human cardiomyocytes^76^. Similarly, lowered body temperature during hibernation of brown bears also results in elevated mRNA level of a CLOCK target gene, *cry2*^77^. Although AS of mammalian *Clock* has been previously identified^78^, whether AS of mammalian *Clock* mediates mRNA expression of its targets in a temperature-sensitive manner remains to be investigated.

We show that increased transcriptional activity of CLK-cold promotes CLK target mRNA expression at cold temperature (Fig. 2). Previous studies suggest that cold-induced intron splicing of *per* and accumulation of *tim*-SC transcripts promote organismal adaptation to cold temperature^13,23,25^. It is possible that the elevated transcriptional activity of CLK due to AS of *clk* transcripts can promote cold adaptation by increasing *per* and *tim*-SC mRNA level. In addition, CLK-cold could regulate cold adaptation through PDF, which can downregulate EYES ABSENT, a seasonal sensor protein that integrates temperature and photoperiodic signals^64,79^. As our lab previously showed that PDF level decreases in flies at 10°C, one potential mechanism by which CLK-cold regulates PDF is to increase the expression of HR38 and/or SR, two CLK targets that can inhibit PDF expression^62^. Moreover, Li et al.^80^ recently identified that DN1a dorsal neurons can modulate locomotor activity and sleep distribution in response to temperature changes. Since both *clk*-long and *clk*-cold are expressed in DN1 neurons^38^, it is possible that AS of *clk* transcripts may contribute to the temperature sensing function of DN1a neurons. Brief inspection of genomic *clk* of several other *Drosophila* species indicates that they also have potential alternative 3’ splice sites that can produce both *clk*-long and *clk*-cold. This indicates the adaptative value of *clk* AS is not limited to *Drosophila melanogaster*, but also in other *Drosophila* species, such as cold-adapted *Drosophila montana* and an agricultural pest, *Drosophila suzukii*. To further understand how flies adapt to cold through AS of *clk*, future studies are necessary to uncover how daily rhythmic transcriptome is altered in the cold.

Within the four amino acids that are spliced out in CLK-cold, we identified S13 as a phosphorylation site adjacent to the bHLH domain that regulates CLK-DNA binding (Fig. 5). We hypothesize that flies expressing the non-phosphorylatable *clk*(S13A) mutant would partially mimic the phenotype of flies under cold conditions. Indeed, CLK binding to DNA is elevated at early morning time in both *clk*(S13A) mutant flies and flies at 10°C (Fig. 2A and B, 5F and G). However, we also observed some discrepancies regarding CLK-DNA binding at later time-points in late day and early evening (Fig. 2A and B, 5F and G) as well as the mRNA levels of CLK targets (Fig. 2D and 4G-I) between these two types of flies. It is important to note that *clk* AS is certainly not the only mechanism mediating temperature responses of the molecular clock. For instance, under cold conditions, *per* and *tim* also undergo AS and alter their repressor activity on CLK, and these mechanisms are not at work in *clk*(S13A) flies under warm temperature.

In addition to interpreting the molecular phenotype of flies at 10°C, the characterization of *clk*(S13A) mutant provided us with new insights into the mechanism by which CLK transcriptional activity is repressed by phosphorylation. Our CLK-ChIP data showed no significant increase of CLK-DNA binding in *clk*(S13A) mutant at late night (ZT23) as compared to *clk*(WT) (Fig. 5F and G), indicating CLK-DNA dissociation is not affected when CLK(S13) phosphorylation is abolished. Rather, CLK-DNA binding is increased in early morning (ZT3) in *clk*(S13A) mutant. We reasoned that upon S13 phosphorylation, CWO outcompetes CLK in *E-box* binding activity, preventing off-DNA CLK from binding back. This is supported by previous findings that PER, likely PER-dependent phosphorylation of CLK, is necessary for CWO to compete with CLK for *E-box* binding^81^. It is somewhat surprising that elevated CLK-DNA binding at ZT3 in *clk*(S13A) mutants does not enhance CLK target gene expression (Fig. 4G-I). This supports the importance of other phosphorylation sites and/or other posttranslational modifications, such as CK2-dependent phosphorylation^49^ and USP8-dependent deubiquitylation^82^, in repressing transcriptional activity of CLK. In conclusion, the primary function of S13 phosphorylation is likely to prevent off-DNA CLK from prematurely associating with DNA in early morning, rather than dissociating CLK from DNA in the evening.

We provide evidence supporting the long-standing hypothesis proposed in 2006 that PER acts as a scaffold to deliver unknown kinases to directly repress transcriptional activity of CLK^28^. DBT was first implicated as the CLK kinase^28,71^, while follow up studies suggest kinase activity of DBT is not required for CLK phosphorylation^32^. Rather, DBT together with PER act as the molecular scaffold for the phosphorylation-dependent repression on CLK. Our lab previously showed that CK1α interacts with PER in both the cytoplasm and the nucleus^50^. Here, we showed that CK1α is a novel kinase of CLK that requires the presence of PER-DBT complex to phosphorylate CLK at S13 (Fig. 3 and 6). As discussed above, CLK S13 phosphorylation inhibits CLK-DNA binding activity. In addition, casein kinase 1 (CK1)-dependent phosphorylation of repressor proteins have been proposed as conserved timing mechanisms in eukaryotic circadian clocks, despite different activator and repressor proteins employed^83^. Together with recent findings in fungi^84^ and mammals^85^, our data further suggest the CK1-scaffolding role of repressor proteins as additional conserved features in regulating eukaryotic circadian clocks.

In summary, we uncovered an interplay between temperature-sensitive AS and phosphorylation in modulating the activity of a master clock transcriptional activator. Many studies have been devoted to investigate the function of AS, as it is prevalent in 42-95% intron-containing genes across species^86–89^. However, adjacent AS sites (<= 18 bps) are usually overlooked due to their perceived minor influence in protein coding, despite its prevalence in transcriptomes among several species^87,90–92^. Our current study provides an example on the importance of adjacent AS sites. Moreover, interplay between AS and phosphorylation have been shown to occur at different levels, including phosphorylation of splicing-related proteins^11,12^, AS of kinases and phosphatases^38,93,94^ and inclusion of cassette exons with phosphorylation sites^95^. Here, we provide an additional mechanism in which AS regulates protein function by removing amino acid(s) as substrate for phosphorylation encoded between adjacent AS sites. Finally, our results also provide novel insights into how the circadian clock maintains its pace in natural conditions with temperature cycles.

## Acknowledgements

We thank Paul Hardin for *attB*-P[acman]-*clk* construct, Carrie Partch for pET22b construct. We thank the Bloomington *Drosophila* Stock Center for providing fly stocks. We thank Alexey A. Tomilov and Gino Cortopassi at UC Davis Drug Discovery Recharge Unit for their technical support on Biolayer Interferometry. The Confocal Microscopy facility was supported by NIH GM122968 to Pamela C. Ronald at UC Davis. We thank Hongtao Zhang for critical reading of the manuscript. Research in the laboratory of JCC is supported by NIH R01 DK124068 and NIH R56 DK124068.

## Author Contributions

J.C.C. and Y.D.C. conceptualized and designed the study. J.C.C. acquired the funding. J.C.C., Y.D.C. and G.K.C. coordinated and supervised the investigation. Y.D.C., G.K.C., S.H., K.C.J., C.D.V., Z.Y.G., V.H.L., H.Z. and C.Z. performed research and analyzed data. Y.D.C., J.C.C., G.K.C., and X.L. contributed to critical interpretation of the data. C.A.T. generated reagents. Y.D.C., X.L. and J.C.C. drafted the manuscript.

## Methods

### Material availability

All unique/stable reagents generated in this study are available from the lead contact without restriction.

### Data and code availability

The codes used in the data analysis and the datasets generated during the current study can be found at Github [https://github.com/ClockLabX/CLK_AS].

### Experimental Models and Subject Details

#### *Drosophila* construct design and transformation

*attB*-P[acman]-*clk* (15.5kb of the genomic sequence beginning ∼8kb upstream and ending ∼2.5kb downstream of *clk* coding region) was kindly provided by Paul Hardin^55^. To introduce a V5 epitope tag in the C-terminus of the *clk* coding region, a 4kb NheI-NotI *clk* fragment was subcloned into pSP72 construct and sequences encoding V5 were introduced in frame by site-directed mutagenesis using Pfu Turbo Cx DNA polymerase (Agilent Technologies, Santa Clara, CA) (See Table S5 for mutagenic primer sequences). The resulting NheI-NotI *clk*-V5 fragment was then used to replace the NheI-NotI *clk* fragment in *attB*-P[acman]-*clk* to generate *attB*-P[acman]-*clk*-V5. PhiC31-mediated transgenesis^96^ was used to generate *w*^1118^; *clk*(WT)-V5. Plasmids were injected into PBac{y[+]-attP-9A}VK00018 fly embryo (Bloomington #9736) (BestGene, Chino Hills, CA)^97^. Transformants were crossed with *w*^1118^; +; *clk*^out^ flies (Bloomington #56754)^55^ to remove endogenous copies of *clk* prior to behavioral and molecular analyses.

To generate flies expressing non-phosphorylatable (Serine (S) to Alanine(A)) or phosphomimetic (S to Aspartic acid (D)) *clk* mutants, a 7kb NheI-SphI *clk* fragment was subcloned into pSP72 plasmid where a NheI site were introduced to multicloning sites. After site-directed mutagenesis and confirmation by Sanger sequencing (GENEWIZ Inc, South Plainfield, NJ), the mutant variants of 7kb *clk* fragments were used to replace the corresponding WT fragment in *attB*-P[acman]-*clk*(WT)-V5. Transgenic flies were generated by Bestgene (Chino Hills, CA) as described above.

### Method Details

#### RNA Extraction, cDNA Synthesis, RT-PCR, and quantitative PCR

RNA was extracted from approximately 30-50µl of fly heads using 3X volume TRI Reagent (Sigma-Aldrich, St. Louis, MO). 1/5 volume of 100% chloroform (Sigma-Aldrich) was added and incubated at room temperature for 10 minutes. Upper aqueous layer was recovered after spinning down at 13,000 rpm for 15 minutes. Same volume of 100% isopropanol was added and incubated at −20°C overnight to precipitate RNA. After spinning down, RNA pellet was washed with 200µl 70% ethanol once, resuspended in 20µl 1X RQ1 buffer (Promega, Madison, WI), and treated with 2µl RQ1 DNase (Promega) at 37°C for 30 minutes prior to the incubation with 2µl RQ1 DNase stop solution (Promega) at 65°C for 10 minutes. cDNA was generated from equal amount of RNA for each sample using Superscript IV (Thermo Fisher Scientific, Waltham, MA). Real-time PCR was performed using SsoAdvanced SYBR green supermix (Bio-Rad, Hercules, CA) in a CFX96 or CFX384 (Bio-Rad). Three technical replicates were performed for each of three biological qPCR replicates.

To confirm expression of *clk* isoforms, RNA was extracted as above. cDNA was generated using gene-specific primer *clk*(698R). PCR was then performed using *clk*(1F) and *clk*(101R) prior to resolving PCR products in 2% TBE gel. Bands were excised, and gel extracted for Sanger sequencing. Sequences for all primers are presented in Table S5.

#### Plasmids for *Drosophila* S2 cell culture

pAc-*clk*(WT)-V5^98^, *per-E-box-Luciferase*^42^, *pCopia Renilla-Luciferase*^99^, pAc-*per*(WT)-V5^100^, pMT-*ck1α*(WT)-c-myc and pMT-*ck1α*(K49R)-FH^50^ (FH denotes 3XFLAG-6XHis) were previously described. pAc-*clk*-cold was generated by deleting 12bp encoding aa 13-16 of CLK-long from pAc-*clk*(WT)-V5 using mutagenic primers in Table S5.

#### *Drosophila* S2 cell culture and transfection

*Drosophila* S2 cells and Schneider’s *Drosophila* medium were obtained from Life Technologies (Carlsbad, CA). S2 cells were grown at 22°C in Schneider’s *Drosophila* medium supplemented with 10% Fetal Bovine Serum (FBS) (VWR, Radnor, PA) and 0.5% Penicillin/streptomycin (Sigma-Aldrich). For all cell culture experiments unless otherwise noted, S2 cells were seeded at 1 × 10^6^ cells/ml in a 6-well plate and transfected using Effectene (Qiagen, Germantown, MD). For coimmunoprecipitation (coIP) assays in Fig. 3B, S2 cells were cotransfected with 0.8µg of pAc-*clk*-V5-His and 0.8µg of pMT-*ck1α*-6Xc-myc, and induced with 500 μM CuSO_4_ immediately after transfection. In control IPs to detect non-specific binding, cells were transfected with either pAc-*clk*-V5-His or pMT-*ck1α*-6Xc-myc, in combination with pMT-FH empty plasmid to balance amount of total transfected plasmids. For mobility shift assay in Fig. 3E and Fig. S3, S2 cells were cotransfected with 0.8µg of pAc-*clk*-V5 in combination with 0.6µg pMT-*ck1α*(WT)-FH, pMT-*ck1α*(K49R)-FH, or pMT-FH. 36 hours following transfection, kinase expression was induced with 500 μM CuSO_4_ for 24 hours and treated with cycloheximide (CHX) (Sigma-Aldrich) (10µg/ml) and MG132 (Sigma-Aldrich) (25µg/ml) for 4 hours. For CLK(S13) phosphorylation detection in Fig. 3H, cells were transfected with 0.8µg of pAc-*clk* (WT or S13A)-V5-His with either 0.6µg of pMT-*ck1α*-FH or pMT-FH. 24 hours following transfection, kinase expression was induced with 500 μM CuSO_4_ for 24 hours. For CLK(S13) phosphorylation detection in Fig. 6D, cells were transfected with 0.8µg of pMT-FH-*clk*(WT), 0.1µg of pMT-*dbt*(K/R), 0.1µg of pMT-*ck1α*-cmyc and pAc-*per*(WT or ϕλ)-NLS-V5 and induced with 500 μM CuSO_4_ immediately after transfection. For CHX chase assay in Fig. S4, S2 cells were transfected with 0.8µg of pAc-*clk*(WT)-V5 and either 0.6µg of pMT-*ck1α*-FH or pMT-FH.

For luciferase reporter assay in Fig. 2C and Fig. 3A, S2 cells were cotransfected with the plasmid combination as indicated: 0.025µg of *per-luc*, 0.025µg of *ren-luc* and 0.002µg of pAc-*clk*(X)-V5 where X is either WT, S13A, S13D or cold isoform. S2 cells were harvested 36 hours after transfection prior to reporter assay. For luciferase reporter assay in Fig. 6, S2 cells were cotransfected with the plasmid combination as indicated: 0.1µg of *per-E-box-Luciferase*, 0.1µg of pCopia *Renilla-luciferase*, 8ng of pAc-*clk*(WT)-V5, 8ng of pMT-*ck1α*-FH and 80ng of pAc-*per*(WT)-V5 or pAc-*per*(WT or Δ)-NLS-V5. Kinase expression was induced with 500 μM CuSO_4_ immediately after transfection and cells were harvested 44 hours after induction.

#### Luciferase reporter assay

Measurements were performed using the Dual-Glo Luciferase Assay System following the instructions of manufacturer (Promega). Two technical replicates were performed for each biological luciferase reporter assay replicates. Three to four biological replicates were performed.

#### Chromatin Immunoprecipitation (ChIP)

CLK-ChIP was performed as described previously^101^. All buffers described below, except ChIP Elution buffer, contain 1X SIGMAFAST EDTA-free protease inhibitor and 0.5 mM PMSF. Briefly, fly head tissues were homogenized using liquid nitrogen chilled mortar and pestle, mixed with Nuclear Extraction buffer (NEB) (10mM Tris-HCl pH 8.0, 0.1mM EGTA pH 8.0, 10mM NaCl, 0.5mM EDTA pH 8.0, 1mM DTT, 0.5% Tergitol NP-10, 0.5mM Spermidine, 0.15mM Spermine), and lysed with a glass dounce homogenizer (Wheaton, Millville, NJ). Homogenate was transferred to a 70μm cell strainer (Thermo Fisher Scientific) prior to centrifugation at 300 g for 1 minute at 4°C. Supernatant were centrifuged at 6700 rpm for 10 minutes at 4°C. Pellets were resuspended in NEB buffer prior to centrifugation at 11,500 rpm for 20 minutes at 4°C on a sucrose gradient (1.6M sucrose in NEB and 0.8M sucrose in NEB). Nuclei-containing pellets were fixed with 0.3% formaldehyde in NEB and rotated at room temperature for 10 minutes. Glycine was then added at a final concentration of 0.13mM to quench crosslinking. Samples were centrifuged at 6,500 rpm for 5 minutes at 4°C. Pellets (cross-linked chromatin) were washed twice with NEB and resuspended in Sonication buffer (10mM Tris-HCl pH 7.5, 2mM EDTA pH 8.0, 1% SDS, 0.2% Triton X-100, 0.5mM Spermidine, 0.15mM Spermine). The cross-linked chromatin was sheared by sonicator (Q80023, QSonica, Newton, Connecticut) to roughly 500 base pair fragments. Supernatant (sheared chromatin) was collected after the centrifugation at 10,000 rpm for 10 minutes. 1.5μl of CLK antibodies^69^ were incubated with 25μl of Dynabeads (Thermo Fisher Scientific) in ChIP Wash buffer (50mM Tris-HCl, 1mM EDTA pH 8.0, 1% Triton X-100, 0.1% DOC, 10μg/ml AcBSA (Promega), 100mM KCl in 1X PBS, 150mM NaCl, 5mM EGTA pH 8.0, 0.1% SDS) at 4°C for 2 hours. Following incubation, beads were collected using a magnet stand (Sigma-Aldrich) and incubated with sheared chromatin that were diluted 10-fold with IP buffer (50mM Tris-HCl pH 7.5, 2mM EDTA pH 8.0, 1% Triton X-100, 0.1% DOC, 150mM NaCl, 0.5mM EGTA pH 8.0) at 4°C for 2 hours. Beads were then collected and washed twice with ChIP Wash buffer for 30 minutes at 4°C, once with LiCl Wash buffer (10mM Tris-HCl pH 8.0, 250mM LiCl, 0.5% NP-40, 0.5% DOC, 1mM EDTA pH 8.0) for 30 minutes at 4°C and once with TE buffer (1mM EDTA pH 8.0, 10mM Tris-HCl pH 8.0) for 4 minutes at 4°C. Beads were eluted with ChIP Elution buffer (50mM Tris-HCl pH 8.0, 10mM EDTA pH 8.0, 1% SDS, 1mM DTT, 50mM NaCl, 4U/ml Proteinase K (NEB, Ipswich, MA), 50μg/ml RNase A (Thermo Fisher Scientific) at 37°C for 2 hours and de-crosslinked at 65°C overnight. Finally, DNA was purified by QIAquick PCR Purification Kit (Qiagen) and quantified by real-time qPCR. Primers for *per CRS*, *tim E-box* were described previously^101^. Primers for *pdp1α E-box* and *vri E-box* are in Table S5. The average of ChIP signals for two intergenic regions, one on chromosome 2R (see Table S5) and one on the X chromosome^101^, was used for non-specific background deduction unless otherwise noted. Three technical replicates were performed for each biological ChIP replicate. At least three biological ChIP replicates were performed.

#### Coimmunoprecipitation experiments in *Drosophila* S2 cells

CoIP experiments were performed as described previously^50^ with the following modifications. *Drosophila* S2 cells were harvested 40 hours after transfection, washed once with 1X PBS and lysed with modified RIPA (20mM Tris-HCl pH 7.5, 150mM NaCl, 10% glycerol, 1% Triton X-100, 0.4% sodium deoxycholate, 0.1% SDS) supplemented with 1mM EDTA pH 8.0, 25mM NaF, 0.5mM PMSF, and SIGMAFAST EDTA-free Protease inhibitor tablet (Sigma-Aldrich). Proteins were incubated with 20μl α-V5 or α-FLAG M2 resins (Sigma-Aldrich) for 4 hours at 4°C to pull down CLK or CK1α, respectively. Resins were washed three times in 500µl modified RIPA buffer at 4°C using end-over-end rotator. Immune complexes were analyzed by Western blotting. Signal intensity of interacting protein was normalized to the intensity of the bait protein. Three biological replicates were performed.

#### Western blotting and antibodies

Western blotting and image analysis were performed as previously described^101^. Upon extraction, protein concentration was measured using Pierce Coomassie Plus Assay Reagents (Thermo Fisher Scientific). 2X SDS sample buffer was added and the mixture boiled at 95°C for 5 minutes. Equal amounts of proteins were resolved by polyacrylamide-SDS gel electrophoresis (PAGE) and transferred to nitrocellulose membrane (Bio-Rad) using Semi-Dry Transfer Cell (Bio-Rad). Membranes were incubated in 5% Blocking Buffer (Bio-Rad) for 40 minutes, incubated with primary antibodies for 16-20 hours. Blots were then washed with 1X TBST for 1 hour, incubated with secondary antibodies for 1 hour, and washed again prior to treatment of Clarity chemiluminescence ECL reagent (Bio-Rad). The following percentage of polyacrylamide-SDS gel were used: 8% for CLK and PER, 10% for HSP70, and 12% for CK1α.

Primary antibodies: α-CLK at 1:2000, α-pS13 at 1:1000, α-V5 (Thermo Fisher Scientific) at 1:1000 for CLK-V5, α-cmyc (Sigma-Aldrich) at 1:2000 for CK1α-cmyc, α-FLAG (Sigma-Aldrich) at 1:7000 for CK1α-FLAG and α-HSP70 (Sigma-Aldrich) at 1:10000. Secondary antibodies conjugated with HRP were added as follows: α-guinea pig IgG (Sigma-Aldrich) at 1:2000 for α-CLK, α-rabbit IgG (GE Healthcare) at 1:1000 for α-pS13, α-mouse IgG (Sigma-Aldrich) at 1:1000 for α-V5 detection, 1:2000 for α-c-myc detection, 1:2000 for α-FLAG detection and 1:10000 for α-HSP70 detection.

#### Phos-Tag gel electrophoresis and Western blotting

*Drosophila* S2 cells were lysed with extraction buffer 2 (EB2) (20mM Hepes pH 7.5, 100mM KCl, 5% glycerol, 1mM DTT, 0.1% Triton X-100, 25mM NaF, 0.5mM PMSF, 10 µg/ml Aprotinin, 5µg/ml Leupeptin, 1µg/ml Pepstatin A). Protein extracts were resolved using 5% SDS-PAGE cast with 10µM of Phos-Tag (Wako, Richmond, VA). Once resolved, gels were incubated for 10 minutes with gentle agitation first in transfer buffer (48mM Tris, 39mM Glycine, 20% Methanol, 0.000375% SDS) containing 1mM EDTA pH 8.0 followed by transfer buffer without EDTA. Proteins were then transferred onto PVDF membranes (Bio-Rad) and visualized by Western blotting. Three biological replicates were performed.

#### Identification of CLK phosphorylation sites from *Drosophila* S2 cells

To generate stable *Drosophila* S2 cell lines for the identification of CLK phosphorylation sites, 1µg of pMT-FH-*clk* and 1µg of pMT-*ck1α*(WT)-6Xc-myc or pMT-*ck1α*(K49R)-6Xc-myc in combination with 1µg of pCoHygro plasmid expressing hygromycin resistance were used for transfection using Effectene (Qiagen). *ck1α*(K49R)-6Xc-myc encodes a kinase dead variant of the kinase. Stable cell lines were established by selection with Schneider’s *Drosophila* medium supplemented with 300µg/ml hygromycin (Roche, Palo Alto, CA).

*Drosophila* S2 cells were harvested by centrifuging at 4,000 rpm for 10 minutes at 4°C. Supernatant was removed and then the cell pellet was washed once with 15ml of 50mM Hepes (pH 7.6). Cells were homogenized in lysis buffer (20mM Hepes pH 7.6, 5% glycerol, 350mM NaCl, 0.1% Triton X-100, 1mM DTT, 1mM MgCl_2_, 0.5mM EDTA pH 8.0, 25mM NaF), supplemented with Complete EDTA-free Protease inhibitor cocktail (Sigma-Aldrich), and PhosSTOP (Roche), by using a 40 ml loose dounce homogenizer (Wheaton). Lysed cells were nutated at 4°C for 30 minutes and then centrifuged at 15,000 rpm for 15 minutes at 4°C. Immunoprecipitation was performed at 4°C overnight with 120μl α-FLAG M2 beads (Sigma-Aldrich) followed by two 10-minute washes using lysis buffer without EDTA, DTT, and PhosSTOP. Bound proteins were eluted with equal bead volume (120µl) of elution buffer (30% glycerol, 3% SDS, 6 mM EDTA pH 8.0, 150 mM Tris pH 6.8) at 95°C for 4 minutes. Eluted proteins were then reduced with 20 mM DTT at 65°C for 20 minutes followed by alkylation at room temperature for 20 minutes with 100 mM Ioacetamide. Proteins were then analyzed by Coomassie staining on a 12% SDS-PAGE gel and CLK containing band was excised for mass spectrometry analysis as described in Chiu et al.^52^.

#### Maxquant and Skyline analysis

Mass spectrometric data were processed with MaxQuant^102^ version 1.6.1.0. MS/MS spectra were searched against the complete Uniprot *Drosophila melanogaster* protein database using the built-in Andromeda peptide search engine^103^ with trypsin designated as the digestion enzyme and two missed cleavages were allowed. Oxidation, N-terminal acetylation, phosphorylation, and deamination of asparagine and glutamine were selected as variable modifications. Carbamidomethylation of cysteine was selected as fixed modification. For all other parameters, MaxQuant default values were selected. Briefly, peptide tolerance for the initial and main search of Andromeda were specified at 20 ppm and 4.5 ppm respectively. For identification, an FDR of 0.01 was selected for peptide spectrum matches (PSM) and protein matches. MaxQuant output data was further processed using Skyline^104^ version 4.1.0.11796. For spectral library building, a cut-off score of 0.95 was selected. For MS1 filtering, precursor with charges of 2, 3, and 4 were considered. For retention time filtering, only scans within 5 minutes of MS/MS identification were selected. Quantification of phosphorylated peptides were performed as area under the curve of each identified peptide.

For quantification of relative phosphopeptide abundance, chromatograms of peptides, shown in Lam^57^, were extracted using Skyline version 4.1.0.11796.

#### Generating CLK(S13) phosphospecific antibodies

Phosphospecific antibodies were generated by PhosphoSolutions (Denver, CO). Rabbits were immunized with a 15-amino-acid peptide (amino acid 6-DDKDDTKpSFLCRKSR-amino acid 20); where pS = phosphoserine). The resulting rabbit sera was further affinity-purified using the pS13 phosphopeptide.

#### Detection of CLK pS13 in *Drosophila* S2 cells

Proteins from S2 cells were extracted using EB2 (20mM HEPES pH 7.5, 100mM KCl, 5% Glycerol, 5mM EDTA, 0.1% Triton X-100, 0.5mM PMSF, 1mM DTT, 10 mg/ml Aprotinin, 5 mg/ml Leupeptin, 1 mg/ml Pepstatin) supplemented with 1X PhosSTOP (Roche) and 25mM NaF. Immunoprecipitation to enrich for CLK proteins was performed as described previously^52^ using 20 μl of α-V5 resin per IP reaction. CLK pS13 is then detected by Western blots using CLK(pS13) phosphospecific antibody.

#### Locomotor activity assay

Daily locomotor activity rhythms in male flies were assayed using the *Drosophila* Activity Monitoring System (DAMS, TriKinetics, Waltham, MA) as described previously^105^.

#### Immunofluorescence and confocal imaging

Brain dissections and immunofluorescence staining procedures were performed as described previously^69^. 3-5-day old flies were entrained for 4 days in 12h:12h LD and fixed with 4% paraformaldehyde for 40 minutes at ZT3 and ZT15 on LD4. Brains were washed three times in 1XPBST (0.1% Triton X-100 in PBS), blocked with 10% Normal Goat Serum (Jackson Immunoresearch, West Grove, PA) in PBST for 90 minutes and incubated with primary antibodies two nights. Primary antibodies against PDF (C7-C; Developmental Studies Hybridoma Bank, Iowa City, IA) was used at 1:1000. Brains were then washed and probed with secondary antibodies α-mouse IgG Alexa Fluor 647 (Jackson Immunoresearch, 115-605-003) at 1:1000. Nine to ten fly brains for each genotype at each time-points were dissected and imaged. Representative images are shown. Fiji software^106^ was used for image analysis.

#### Cycloheximide (CHX) chase assay

24 hours following S2 cell transfection, *ck1α* expression was induced for 16 hours prior to treatment with CHX to stop protein synthesis (10µg/ml) (Sigma-Aldrich). Cells were then harvested and lysed with EB2 supplemented with 5mM EDTA pH 8.0 at the times indicated after CHX addition. Protein lysates were analyzed by Western blotting.

#### Cloning, expression and purification of CLK-bHLH

Unless otherwise specified, LB Broth (Lennox) (Sigma-Aldrich) was used as the culturing media for *Escherichia coli*. Cloning was performed in DH5α cells (Thermo Fisher Scientific). The coding sequence corresponding to the bHLH domain of *Drosophila* CLK (aa 1-71) was PCR amplified from pAc-*clk*(WT)-V5 (Kim and Edery, 2006) and introduced into the expression vector pET22b using the restriction sites NdeI and SalI (New England Biolabs). A His_6_-tag-STOP sequence was introduced into the SalI-reverse primer to enable affinity purification via immobilized metal affinity chromatography (IMAC). S13D mutagenesis was performed using mutagenic primers and Pfu Turbo Cx DNA polymerase (Agilent Technologies, Santa Clara, CA) (See Table S5 for mutagenic primer sequences).

For protein expression and purification, BL21(DE3) (Thermo Fisher Scientific) was transformed with the resultant plasmids, pET22b-CLK71-His_6_ and pET22b-CLK71(S13D)-His_6_. Single colonies were picked from each clone and inoculated into 25-ml starter cultures and grown at 37°C until OD_600_ ∼ 0.5 was reached. Then, the cultures were diluted at a 1:100 ratio in 2 500-ml expression cultures each (1L per construct) and grown at 30°C until OD_600_ ∼ 0.5 was reached. Cultures were then transferred to 4°C for 30 minutes, induced with 1 mM isopropyl-β-D-1-thiogalactopyranoside (IPTG), supplemented with 2% glycerol (final concentrations), and grown at 18°C for 16 hours. Cells were harvested by centrifuging at 4°C at 4,000 rpm for 15 minutes (Sorvall) and resuspended in 40 ml lysis buffer (50 mM sodium phosphate, 500 mM NaCl, 10 mM β-mercaptoethanol, pH 8.0) per construct. The resuspended cells were sonicated over ice with a sonicator (Sonic Dismembrator Model E150E, Thermo Fisher Scientific) at 90% amplitude for 10 cycles of 1-min on and 1-min off. The lysates were then treated with 10 μl DNase I (New England Biolabs) and incubated at 4°C for 1 hour on a rotator followed by centrifugation at 13,000 rpm, 4°C for 30 minutes to remove cell debris. The clarified lysates were then filtered through 0.22 μm filters (EMD Millipore, Burlington, MA) and applied onto an nickel(II)-nitrilotriacetic acid (Ni-NTA) IMAC column (Bio-Rad) on a chromatography platform (NGC, Bio-Rad), washed with 82.5 mM imidazole and eluted on a 82.5 mM – 250 mM imidazole gradient. Fractions containing CLK71-His_6_ or CLK71(S13D)-His_6_ were pooled and dialyzed to working buffer (1× PBS supplemented with 10 mM β-mercaptoethanol) using dialysis cassettes (3.5K MWCO, Thermo Fisher Scientific). Concentrations were measured using Coomassie Plus reagent (Thermo Fisher Scientific) using bovine serum albumin (BSA) (Thermo Fisher Scientific) as a standard. The purified constructs were aliquoted, frozen in liquid nitrogen, and stored at −80°C until further use.

#### Size exclusion chromatography

CLK71-His_6_ or CLK71(S13D)-His_6_ constructs were diluted in working buffer to 50 μM. 250 μL of the diluted proteins were loaded onto a Bio-Rad ENrich 70 10 x 300 column and eluted with working buffer. The column was pre-calibrated with 250 μL of 5-fold diluted gel filtration standard (Bio-Rad) and fitting the peak positions of chicken ovalbumin (44 kDa), equine myoglobin (17 kDa), and vitamin B12 (1350 Da), to the formula log_10_ *M_w_* = *aV* + *b*, where *a* and *b* were empirically determined. The molecular weight of the CLK constructs in solution were then estimated from their peak positions in their chromatograms.

#### Biolayer Interferometry (BLI) data acquisition

BLI was performed using the Octet RED384 system (Sartorius, Göttingen, Germany). All steps were performed in Kinetics buffer (1X PBS, 0.02% Tween-20, 0.1% BSA, 0.05% sodium azide, pH 7.4, Sartorius) unless otherwise stated. For the BLI bait, the 21-bp *per* promoter (5’-CCGCCGCTCACGTGGCGAACT-3’) and scrambled (5’-GTACGCTGCAGGCCCCCTGAC-3’, identical GC content) DNA were purchased from IDT (San Diego, CA). In both cases, the forward oligomers were purchased as 5’ biotin conjugates and annealed in-house with their respective unlabeled complementary strands (1 nmol each) at 95°C for 2 minutes and allowed to cool down to room temperature in a heat block, after which both were diluted to 200 nM in autoclaved water. Nonspecific binding between biosensors and the CLK constructs independent of DNA binding were quantified by functionalizing the Octet Streptavidin (SA) biosensors with biotin (200 nM) only.

For the CLK71-His_6_ or CLK71(S13D)-His_6_ constructs, 675 μL of 20 μM proteins in Kinetics buffer were prepared by diluting the dialyzed proteins in PBS and supplementing with 10X Kinetics buffer. Then, serial dilutions were performed at a 2:1 dilution ratio for 13 points, resulting in a protein concentration range of 20 μM to 103 nM. Additional wells were filled with Kinetics buffer to account for baseline drift. To obtain BLI binding kinetics, SA Biosensors (Sartorius) were treated as follows: (1) preconditioning/soaking (60s), (2) biotin-DNA functionalization in autoclaved water (200s), (3) baseline (180s), and (4) binding (300s). Data were collected at a sampling frequency of 5 Hz (0.2 s intervals).

#### BLI data analysis

The raw BLI kinetic trace data were preprocessed on OctetAnalysis (Sartorius) and then exported to MATLAB (Natick, MA) for further analysis using custom codes. The response arising from CLK-DNA binding, ΔΔR, was computed by subtracting individual kinetic traces by means of double subtraction of (i) traces generated from biosensors functionalized with biotin only and (ii) traces generated from DNA-coated biosensors in the absence of CLK-bHLH, as implemented in OctetAnalysis. As we observed negative response during initial binding (*t* = 0.2 s) for some ΔΔR traces when the biosensors were switched between baseline and binding steps after default preprocessing, indicating incomplete inter-step correction of optical artifacts and/or slight changes in buffer composition, we performed additional custom preprocessing in MATLAB by adjusting the baseline for the binding such that the responses would reach exactly 0 nm when extrapolated to *t* = 0 s. Then, the quasi-steady state response, defined as the average ΔΔR in the time window *t* = 290 – 300 s or the last 10 seconds of the binding step, was plotted against initial CLK concentration [CLK]_0_ and fitted to a 4-parameter Hill equation model,

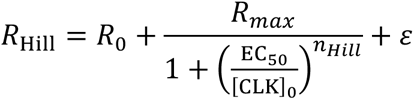

where *R*_0_ is the basal response, *R*_max_ is the maximum change in response elicited by binding, EC_50_ is the half maximal effective concentration, *n*_Hill_ is the Hill coefficient, and *ε* is the error term. To determine if the data could be explained by nonspecific interactions alone, we also fitted the same data to the noise model,

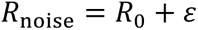

The model-dependent parameters were fitted for each CLK construct-DNA bait pair using the MATLAB function `fitnlm` with *R*_max_, EC_50_, and *n*_Hill_ constrained to nonnegative values via logarithmic transform. The 95% confidence intervals were estimated by the Wald method (function `coefCI`). The corrected Akaike information criteria, AICc, was used to determine whether the data was better explained by specific (Hill model) or nonspecific (Noise model) binding.

#### Statistical analysis

RAIN^40^ and CircaCompare^59^ test were performed in R. Other statistical analyses were performed using GraphPad Prism 10.0 (GraphPad Software, La Jolla, California). Two-tailed Student’s t test were performed if only two groups were compared. ANOVA was performed if there are more than two groups compared. One-Way ANOVA and Dunnett post hoc test were chosen if there are one independent variables, whereas Two-Way ANOVA and Šídák’s post hoc test were chosen if there are two independent variables. Asterisks indicate significant differences in mean values between genotypes or conditions at indicated time-points.

---

**Fig. S1.**
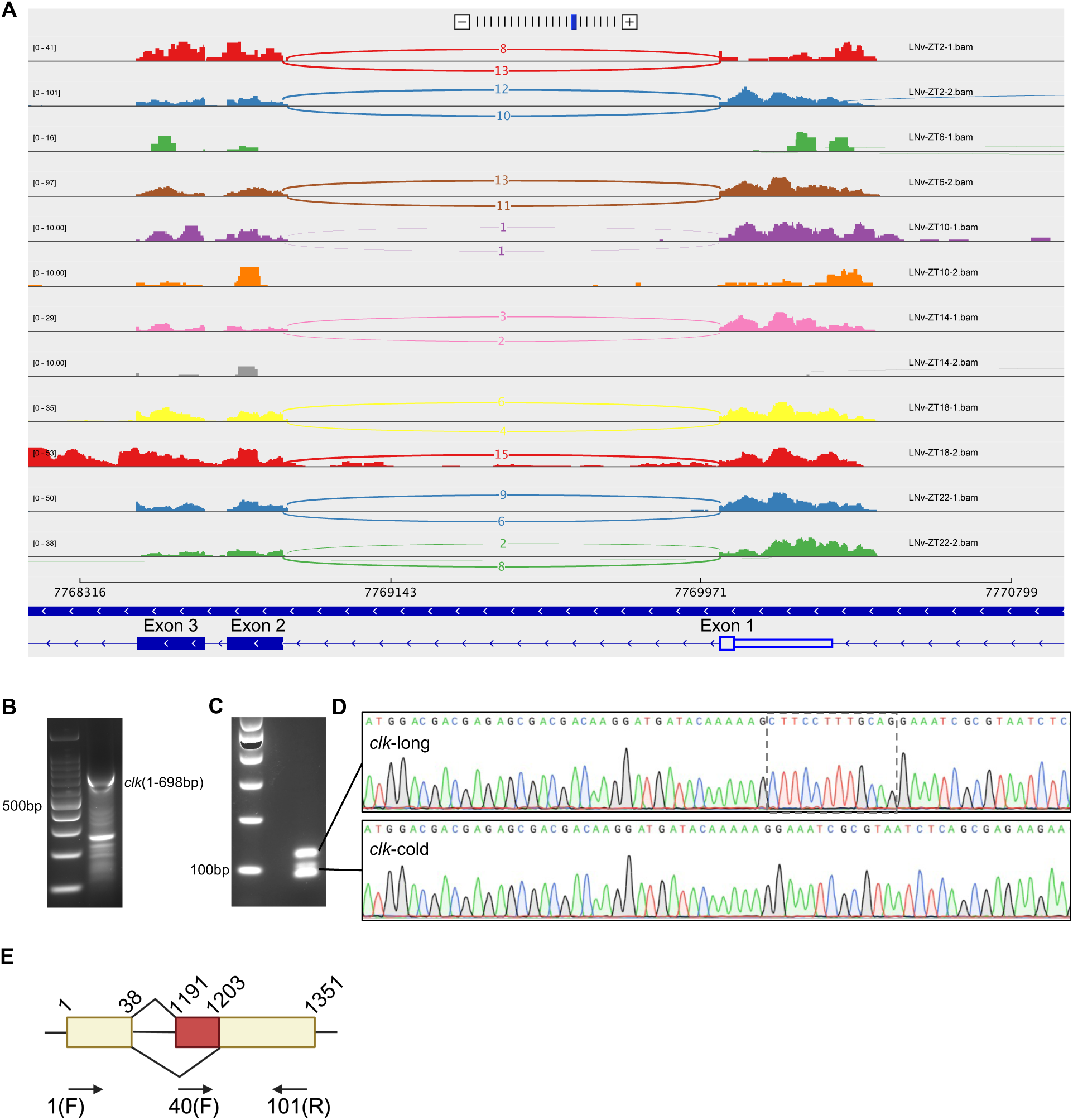
Expression of *clk*-cold isoform in heads of *w*^1118^ flies. **A.** RNA-seq data tracks from Wang et al^1^. are shown to illustrate alternative splicing of *clk* transcripts at exon 2 in small LNv_s_ circadian neurons, with arcs showing splice junctions and the number of unique-mapped RNA-seq reads mapped to the junction across the arc. The orientation of the transcript is indicated by blue arrows at the bottom. **B.** Agarose gel showing reverse transcription product from heads of *w*^1118^ flies collected at ZT0 on LD3 using *clk*(698R) gene-specific primer, amplified by PCR using *clk*(1F) and *clk*(698R) primers (see Table S5 for sequences of primers). **C.** Agarose gel showing *clk*-long and *clk*-cold isoforms amplified by PCR using *clk*(1F)-*clk*(101R) primers with gel extract from **B**. **D.** Chromatogram that confirms the isolation of each isoform. **E.** Schematic representation of primers to measure the relative levels of two *clk* transcripts that vary at the alternative 3’ splice site at exon 2. Exon-intron organization of *clk* and the relative positions of the three primers used in this study for nested RT-qPCR. Two *clk* transcripts quantified by qPCR via either retention (*clk*-long) or removal (*clk*-cold) of the 12bp of exon 2 (indicated in red).

**Fig. S2.**
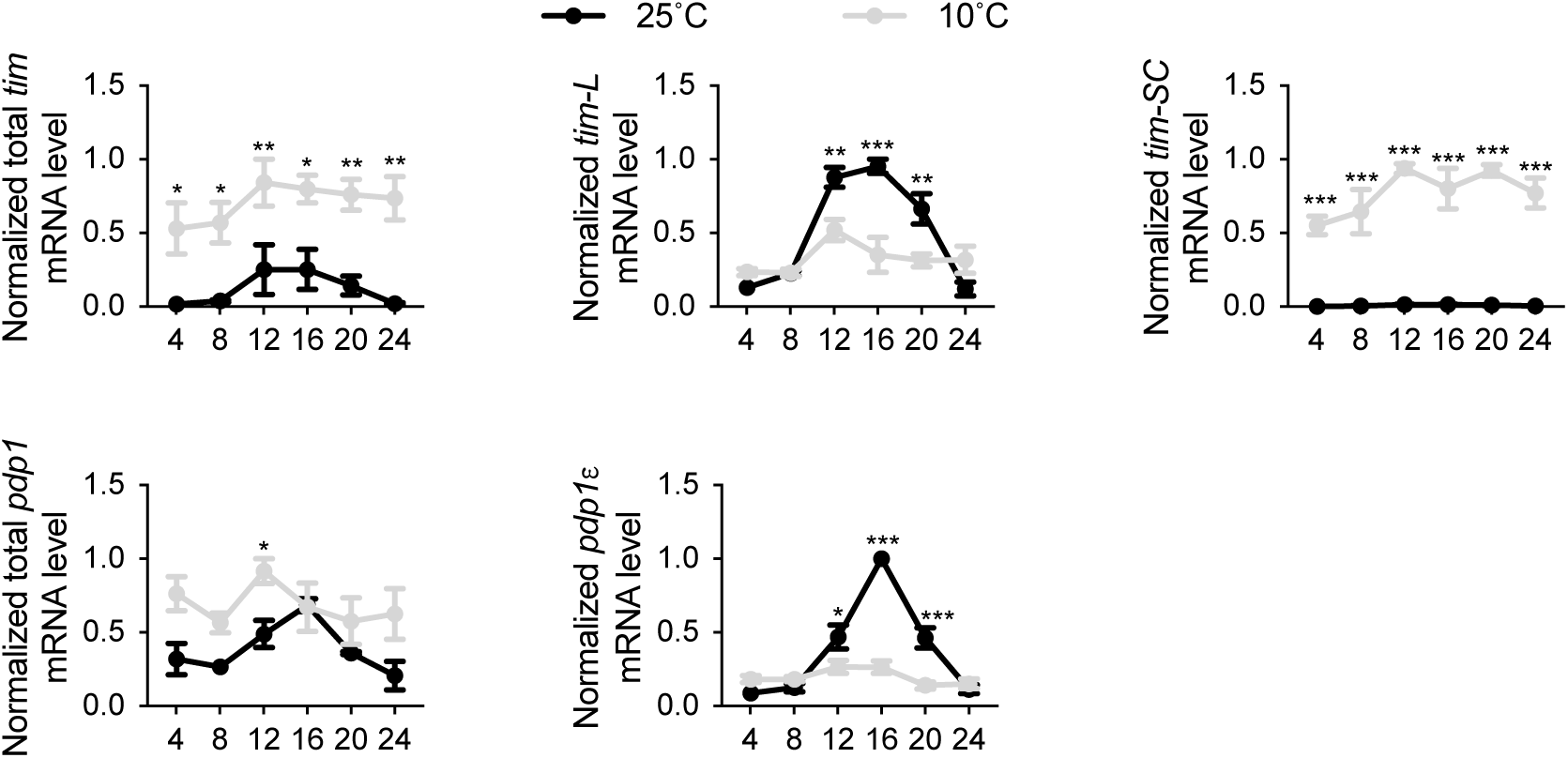
Cold temperature promotes mRNA expression of CLK targets. Daily steady state mRNA expression of CLK targets (*tim* and *pdp1*) in heads of *w*^1118^ flies. *tim*-L, *tim*-SC, *pdp1α* isoforms were analyzed with isoform-specific primers. Flies were entrained in 12h:12h LD and collected on LD3 at the indicated temperatures and time-points (ZT) (n=3). Error bars indicate ± SEM, ***p<0.001, **p< 0.01, *p< 0.05, Two-Way ANOVA and Šídák’s post hoc test.

**Fig. S3.**
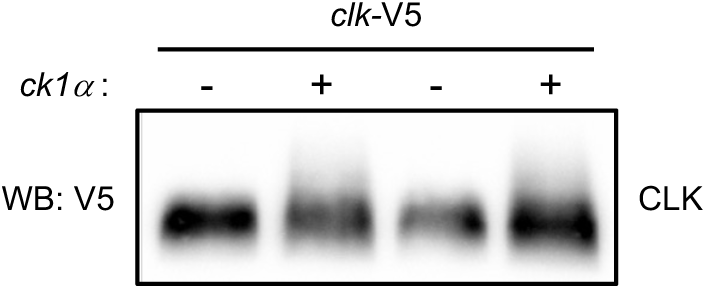
CK1α induces detectable but mild mobility shift of CLK on regular SDS-PAGE gel. *Drosophila* S2 cells were transfected with pAc-*clk*-V5 in combination with either pMT-*ck1α*-3XFLAG-6XHIS or pMT-3XFLAG-6XHIS empty plasmid. Protein extracts were analyzed on regular SDS-PAGE gel followed by western blotting with α-V5. Blots of two biological replicates were shown on the same gel.

**Fig. S4.**
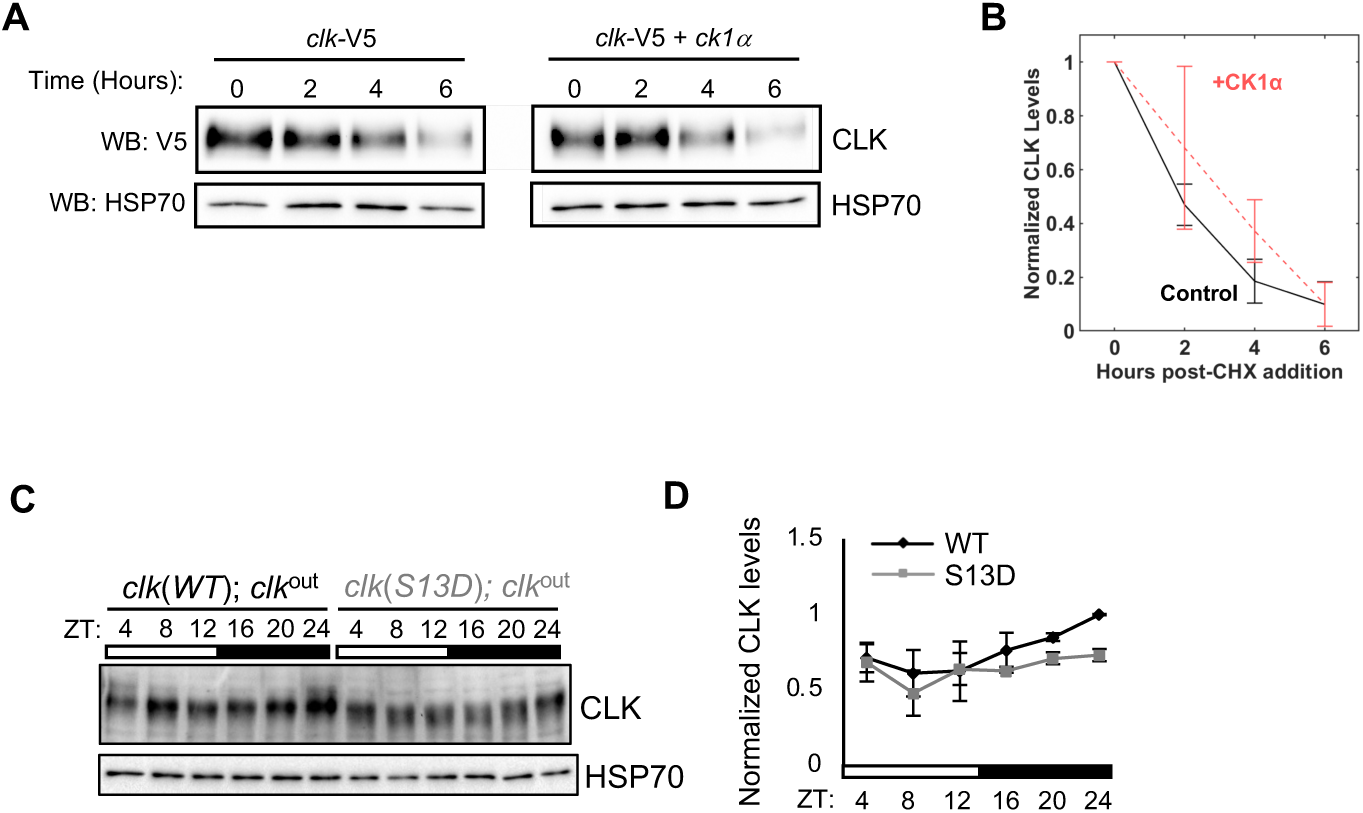
CK1α does not regulate CLK stability. **A.** *Drosophila* S2 cells were cotransfected with pAc-*clk*-V5-His in combination with either pMT-*ck1α*-FH or pMT-FH empty plasmid. Following a 24-hour incubation period, kinase expression was induced with CuSO_4_. Cycloheximide (CHX) was added 16 hours after kinase induction and cells were harvested for protein extractions at the indicated times after addition of CHX. Proteins were visualized by Western blotting and detected with α-V5. α-HSP70 was used to indicate equal loading and for normalization. **B.** Quantification of CLK in **A**. Error bars indicate ± S.E.M (n=2). **C.** Western blots comparing CLK protein profiles in heads of *clk*(WT) and *clk*(S13D) entrained in 12h:12h LD and collected at indicated time-points on LD3. ⍺-HSP70 was used to indicate equal loading and for normalization. **D.** Quantification of CLK in **C** (n=3). Error bars indicate ± SEM, **p< 0.01, Two-Way ANOVA and Šídák’s post hoc test.

**Fig. S5.**
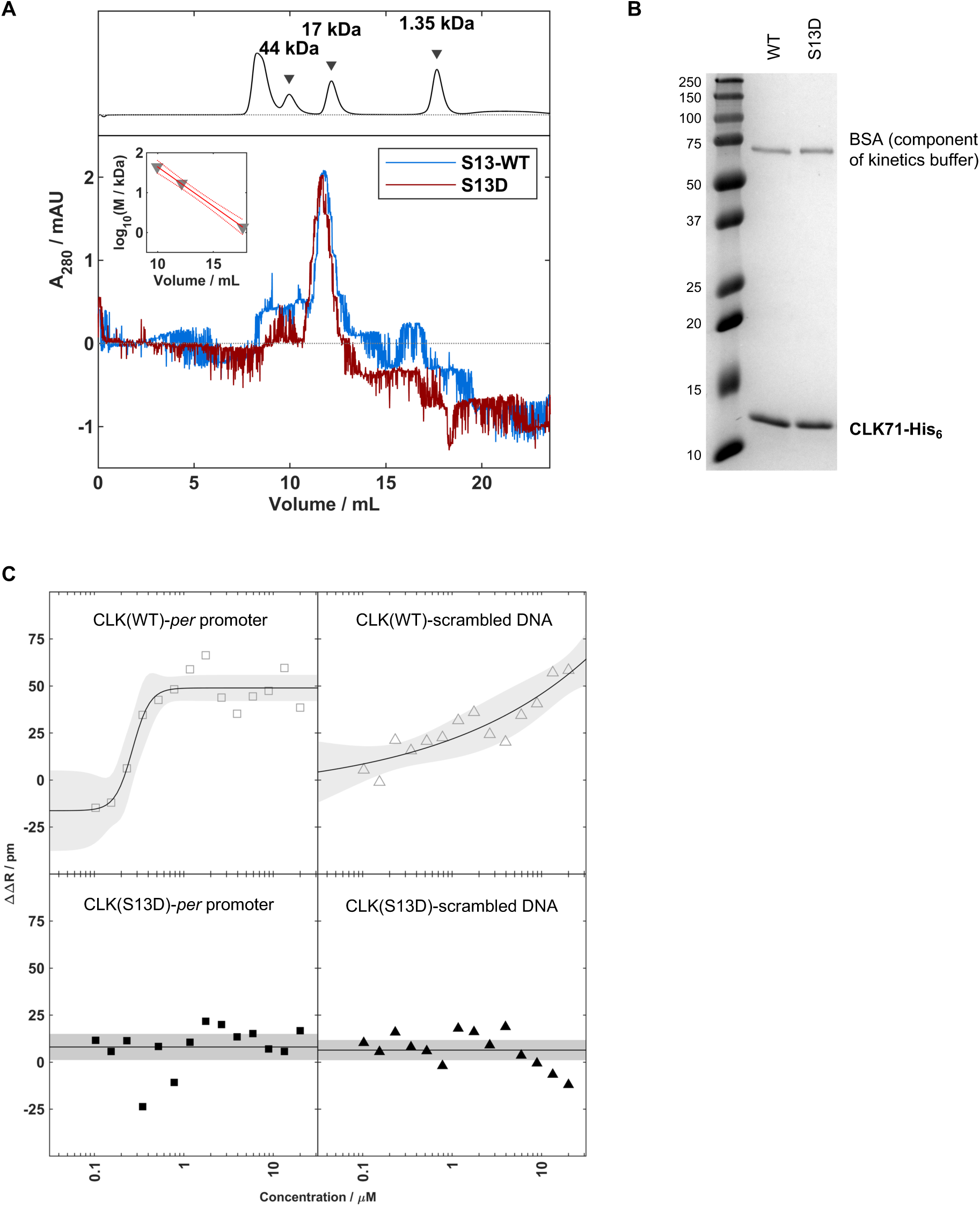
Additional characterization of CLK-bHLH constructs. **A.** Size exclusion chromatography of CLK71-His6 constructs suggests that both constructs are homodimeric (S13-WT, blue, 19.7 kDa, 95% CI = [15.3, 27.1]; S13D, dark red, 20.4, 95% CI = [15.3, 27.1]) assuming globular structure. The calibration standard with the peaks of chicken ovalbumin (44 kDa), equine myoglobin (17 kDa), and vitamin B12 (1350 Da) are shown above, whereas the standard curve is shown in the inset. **B.** Coomassie-stained 16% SDS-PAGE of constructs in 1X Kinetics buffer (containing 0.1% BSA) used in Biolayer Interferometry. **C**. Quasi-steady state signal response of CLK-bHLH-DNA binding in the presence (black, filled) and absence (grey, hollow) of the phosphomimetic S13D mutation are plotted separately for the 21-bp *per* promoter DNA sequence (squares) or a scrambled sequence with identical GC content (triangles). Solid lines and shaded areas show fits and 95% prediction interval to the 4-parameter Hill equation for CLK-bHLH-WT and nonbinding baseline for CLK-bHLH-S13D.

**Table S1.**
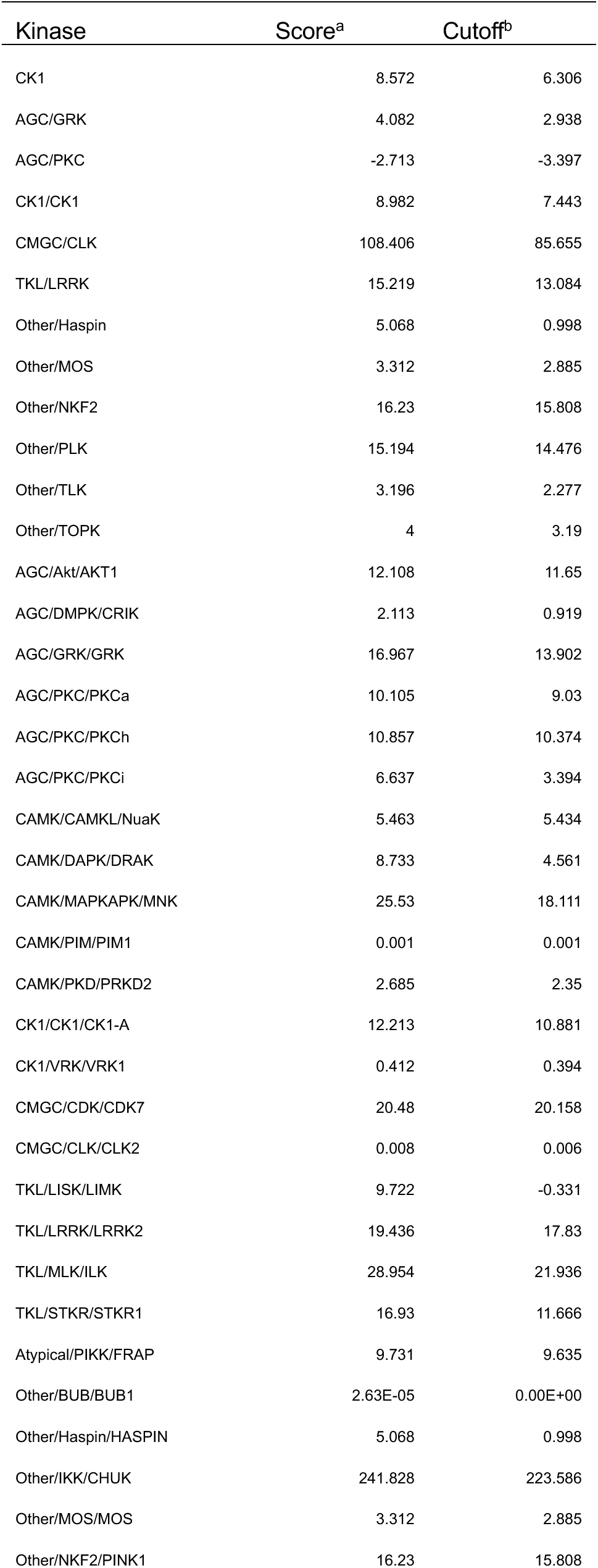
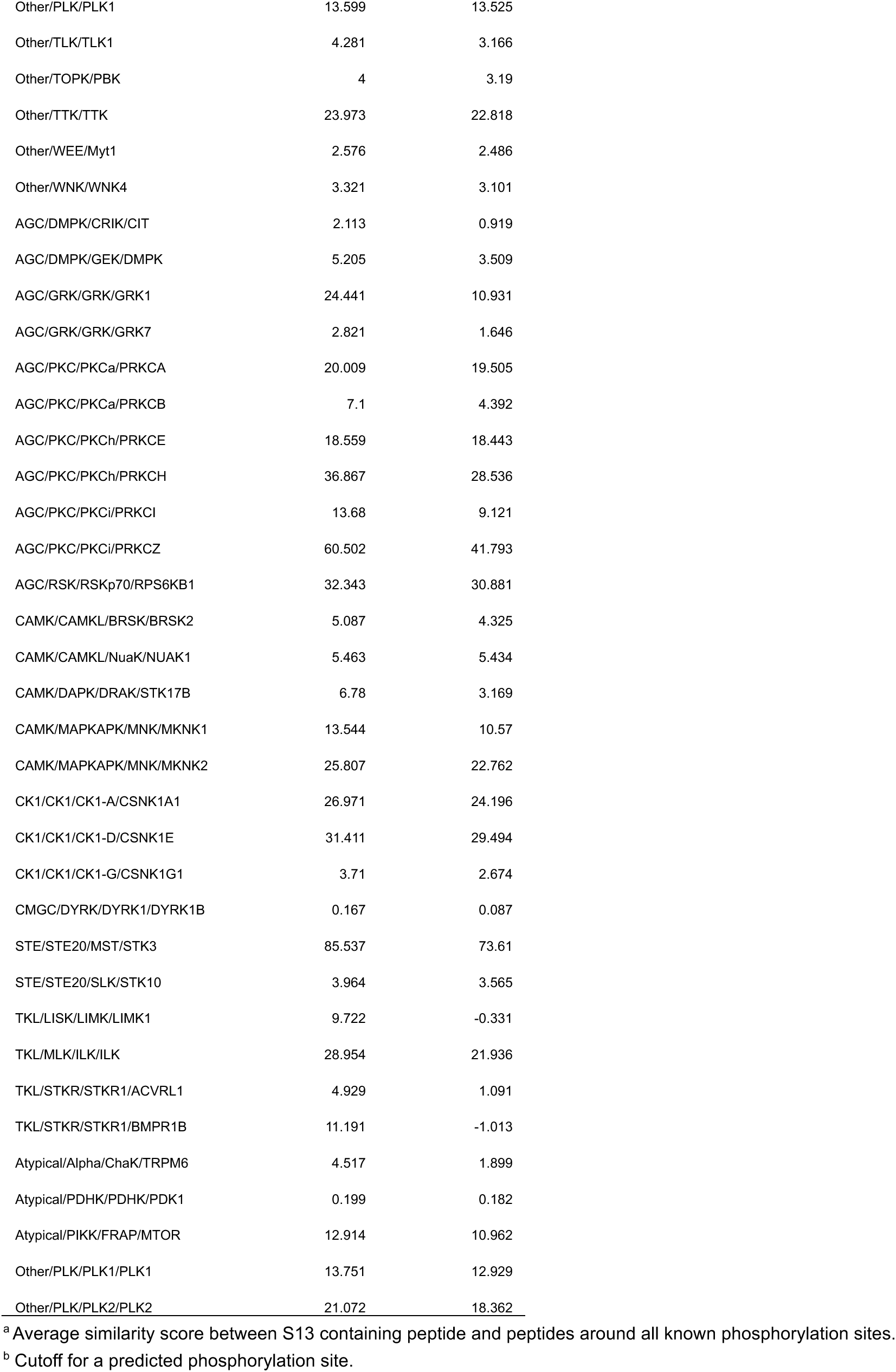
Potential kinases that phosphorylate CLK(S13) predicted by GPS5.0^2^.

**Table S2.**
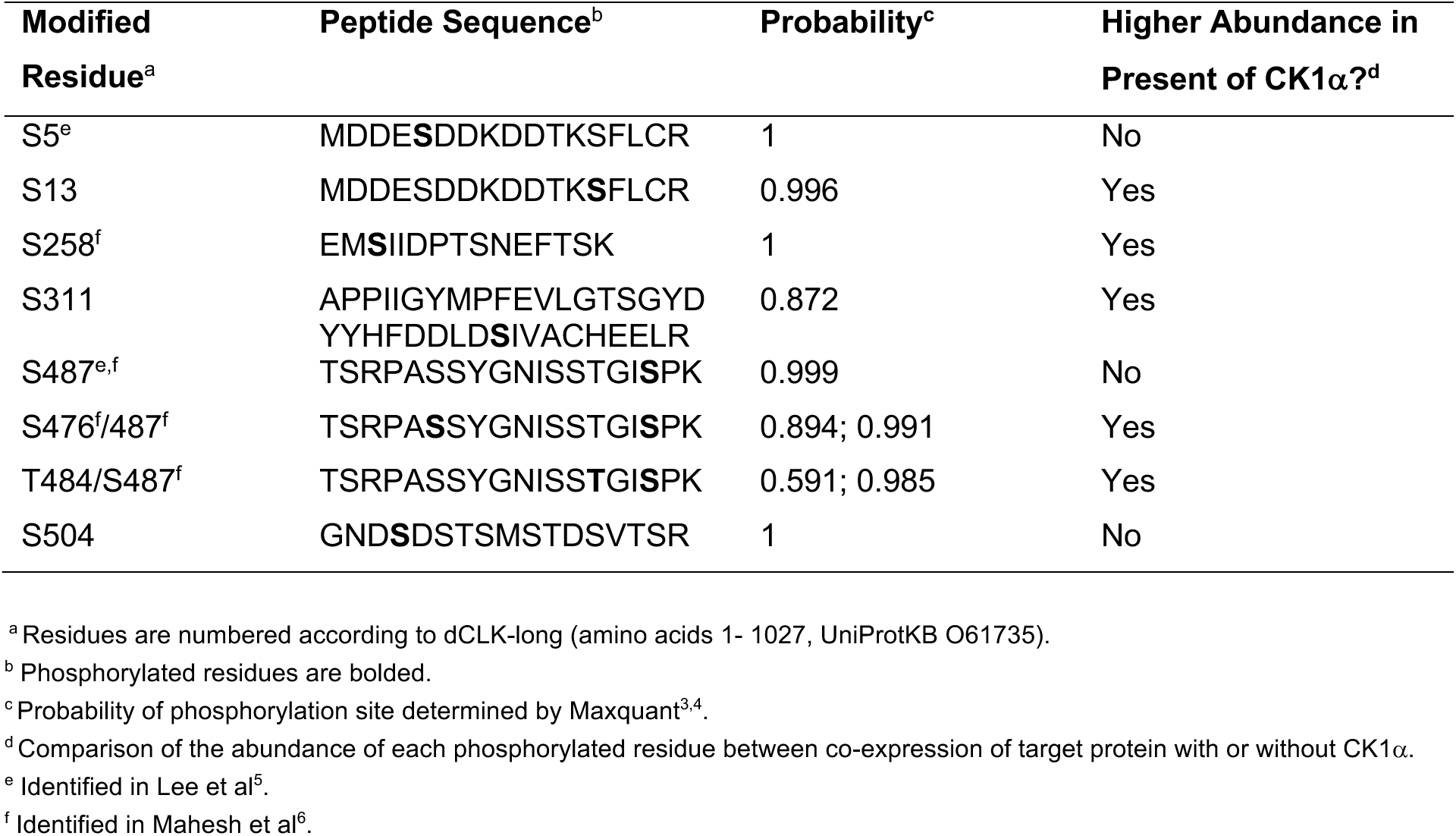
Identification of Ck1α-dependent phosphorylation sites in *Drosophila* head tissues.

**Table S3.** Daily locomotor activity rhythms of *clk* mutants.

| Genotype | Period (h)<br>(mean ± SEM) | Power<sup>a</sup> | Rhythmicity<br>(%)<sup>b</sup> | No. of<br>flies<br>tested | No of flies<br>surviving<sup>c</sup> |
| --- | --- | --- | --- | --- | --- |
| *w; +; clk<sup>out</sup>* | AR<sup>d</sup> | ND<sup>e</sup> | 0 | 32 | 31 |
| *w; clk(WT); clk<sup>out</sup>* | 24.2 ± 0.10 | 68.7 | 86.2 | 63 | 58 |
| *w; clk(WT)/Cyo; clk<sup>out</sup>* | 24.8 ± 0.12 | 59.9 | 61.7 | 64 | 60 |
| *w; clk(S13A)/Cyo; clk<sup>out</sup>* | 25.9 ± 0.13 | 42.0 | 30.4 | 27 | 23 |
| *w; clk(S13D)/Cyo; clk<sup>out</sup>* | 23.5 ± 0.20 | 35.5 | 11.1 | 37 | 36 |
| *w; clk(S13D); clk<sup>out</sup>* | 25.4 ± 0.39 | 50.6 | 39.3 | 32 | 28 |

<sup>a</sup> Measures the strength or amplitude of the locomotor activity rhythm (in arbitrary units)
<sup>b</sup> Percentage of flies that are rhythmic
<sup>c</sup> Number of flies that survived until the end of the experiment
<sup>d</sup> AR denotes Arrhythmic
<sup>e</sup> ND denotes Not Determined

**Table S4.** Rhythmic parameters of mRNA analysis.

| Target | WT mesor | S13D mesor | P-value for mesor difference | WT amplitude | S13D amplitude | P-value for amplitude difference | WT peak time (ZT) | S13D peak time (ZT) | P-value for difference in phase |
| --- | --- | --- | --- | --- | --- | --- | --- | --- | --- |
| *per* | 0.617 | 0.436 | *** | 0.310 | 0.174 | * | 13.215 | 15.256 | * |
| *tim* | 0.521 | 0.208 | *** | 0.395 | 0.158 | *** | 15.743 | 15.906 | 0.755 |
| *vri* | 0.500 | 0.241 | *** | 0.402 | 0.131 | *** | 13.967 | 14.385 | 0.683 |
| Target | WT mesor | S13A mesor | P-value for mesor difference | WT amplitude | S13A amplitude | P-value for amplitude difference | WT peak time (ZT) | S13A peak time (ZT) | P-value for difference in phase |
| *per* | 0.525 | 0.367 | * | 0.390 | 0.173 | ** | 14.173 | 12.039 | 0.079 |
| *tim* | 0.503 | 0.288 | *** | 0.417 | 0.239 | *** | 15.492 | 15.153 | 0.558 |
| *vri* | 0.496 | 0.220 | *** | 0.467 | 0.156 | *** | 13.895 | 12.350 | 0.091 |

**Table S5:**
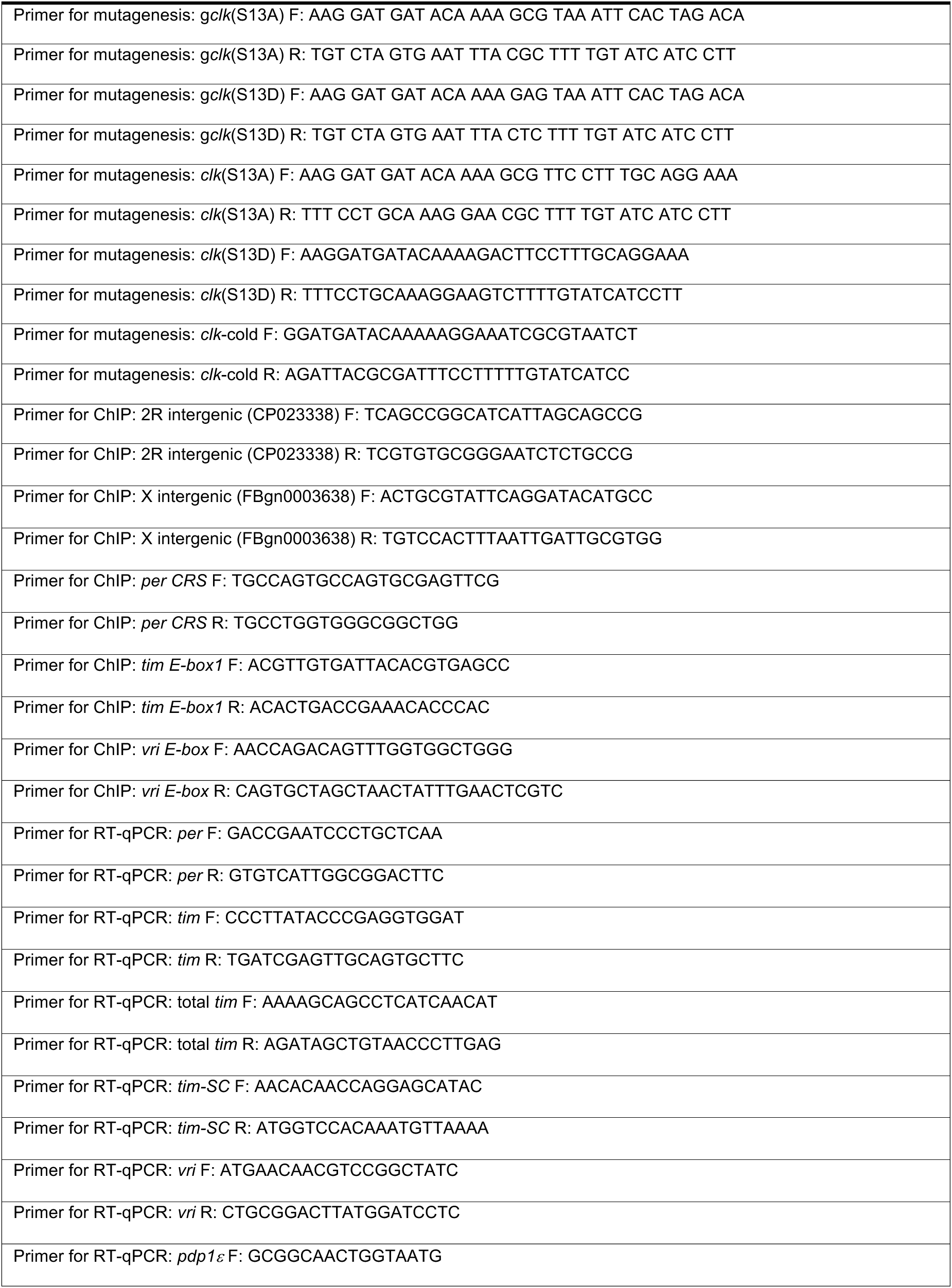
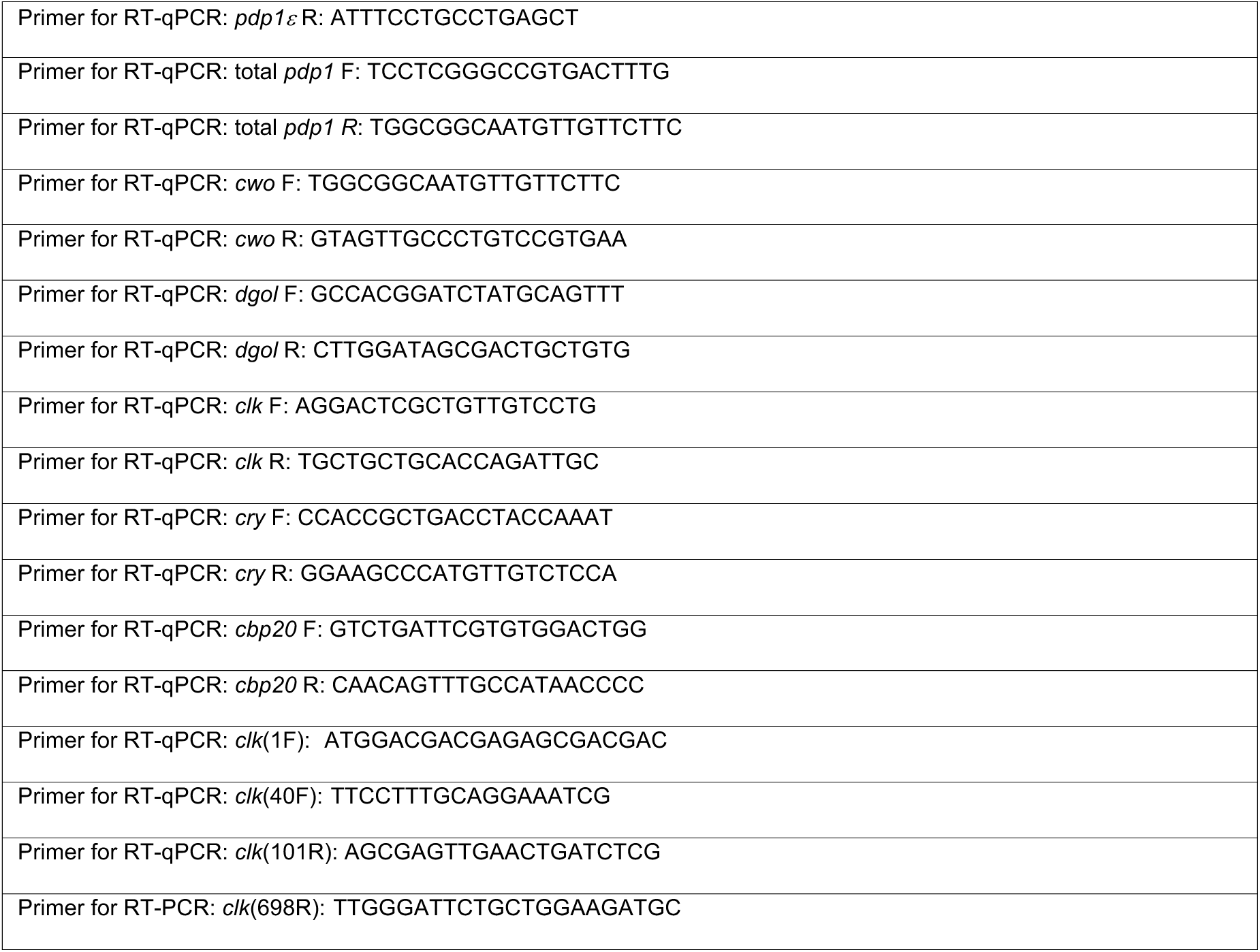
Primers for PCR mutagenesis, ChIP analysis and RT-qPCR analysis.

